# Molecular mechanism of GTP binding- and dimerization-induced enhancement of Sar1-mediated membrane remodeling

**DOI:** 10.1101/2022.07.20.500836

**Authors:** Sanjoy Paul, Anjon Audhya, Qiang Cui

## Abstract

The Sar1 GTPase initiates Coat Protein II (COPII)-mediated protein transport by generating membrane curvature at subdomains on the Endoplasmic Reticulum, where it is activated by the guanine nucleotide exchange factor (GEF) Sec12. Crystal structures of GDP- and GTP-bound forms of Sar1 suggest that it undergoes a conformational switch in which GTP binding enhances the exposure of an amino-terminal amphipathic helix necessary for efficient membrane penetration. However, key residues in the amino-terminus were not resolved in crystal structures, and experimental studies have suggested that the amino-terminus of Sar1 is solvent-exposed in the absence of membrane, even in the GDP-bound state. Therefore, the molecular mechanism by which GTP binding activates the membrane remodeling activity of Sar1 remains unclear. Using atomistic molecular dynamics simulations, we compare the membrane binding and curvature generation activities of Sar1 in its GDP- and GTP-bound states. We show that in the GTP-bound state, Sar1 inserts into the membrane with its complete (residues 1-23) amphipathic amino-terminal helix, while Sar1-GDP binds to the membrane only through its first 12 residues. Such differential membrane binding modes translate into significant differences in the protein volume inserted into the membrane. As a result, Sar1-GTP generates positive membrane curvature 10-20 times higher than Sar1-GDP. Dimerization of the GTP-bound form of Sar1 further amplifies curvature generation. Taken together, our results present a detailed molecular mechanism for how the nucleotide-bound state of Sar1 regulates its membrane binding and remodeling activities in a concentration dependent manner, paving the way toward a better understanding COPII-mediated membrane transport.

**Significance Statement:** Amphipathic helices play established roles in penetrating lipid bilayers to produce local membrane curvature. The degree of curvature generated has been suggested to depend on the penetration depth of the amphipathic helix, its amino acid sequence, and the protein volume inserted into the membrane. However, the relative contributions of each of these factors in membrane bending remain unclear. Using the Sar1 protein monomer bound to different nucleotides (GDP vs. GTP) and its GTP-bound dimeric form as examples, we explicitly show that while the precise amino acid sequence and insertion depth of the amphipathic helix are relevant, the generated membrane curvature is most correlated with the volume of protein insertion into the membrane.

---

**T**ransport of newly synthesized proteins within membrane-enclosed vesicular carriers represents a crucial step in the early secretory pathway of all eukaryotic cells. These carriers are often coated with cytosolic proteins that possess the ability to remodel lipid bilayers. In particular, Coat Protein complex II (COPII) has been demonstrated to direct the trafficking of biosynthetic cargoes that are exported from the Endoplasmic Reticulum (ER).(1) The COPII complex consists of the small GTPase Sar1, the inner coat complex (Sec23-Sec24) and the outer coat complex (Sec13-Sec31).(2) Current evidence suggests that Sar1 initially binds to the ER membrane upon GTP binding, which is facilitated by the guanine nucleotide exchange factor (GEF) Sec12.(3) In the presence of GTP, Sar1 initiates ER membrane tubulation, leveraging an amino-terminal amphipathic helix that penetrates the bilayer.(4) Next, Sec23-Sec24 binds to activated Sar1-GTP, forming a lattice-like assembly.(5) Sec23-Sec24 also plays a pivotal role in sorting cargo molecules for incorporation into the transport carrier.(6, 7) The inner coat complex enables recruitment of Sec13-Sec31 heterotetramers, which form a cage, completing COPII coat formation. While this general model is widely accepted, molecular details remain absent. In particular, the molecular mechanism by which Sar1 generates membrane curvature in a nucleotide-dependent manner remains unclear.

Crystal structures of Sar1 have been solved with either GDP(8) or GNP, a non-hydrolyzable analogue of GTP.(9) While the entire amino-terminal helix (residues 1-23) was not resolved in the GNP-(GTP) bound state, part of the helix (residues 12-23) was observed in the GDP-bound state to retract into a surface pocket formed partly by the loop connecting the *β*2-*β*3 hairpin (Fig. 1A, B). Therefore, it was suggested that GTP binding triggers the exposure of the amino-terminal amphipathic helix, which can anchor, penetrate and bend the membrane. This conformational switch mechanism is consistent with that of ARF family GTPases, for which membrane binding and coat assembly are also regulated through GDP/GTP exchange.(10) However, unlike the case of Sar1, myristoylation in the amino-terminal membrane anchor of ARF family proteins plays a crucial role in membrane binding and nucleotide exchange.(11, 12) GTP binding on ARF induces structural changes in the switch 1 and 2 regions, causing a 2-residue register shift of the interswitch region, blocking binding to the myristoylated amino-terminus and thereby facilitating insertion into the membrane.(13)

**Fig. 1.**
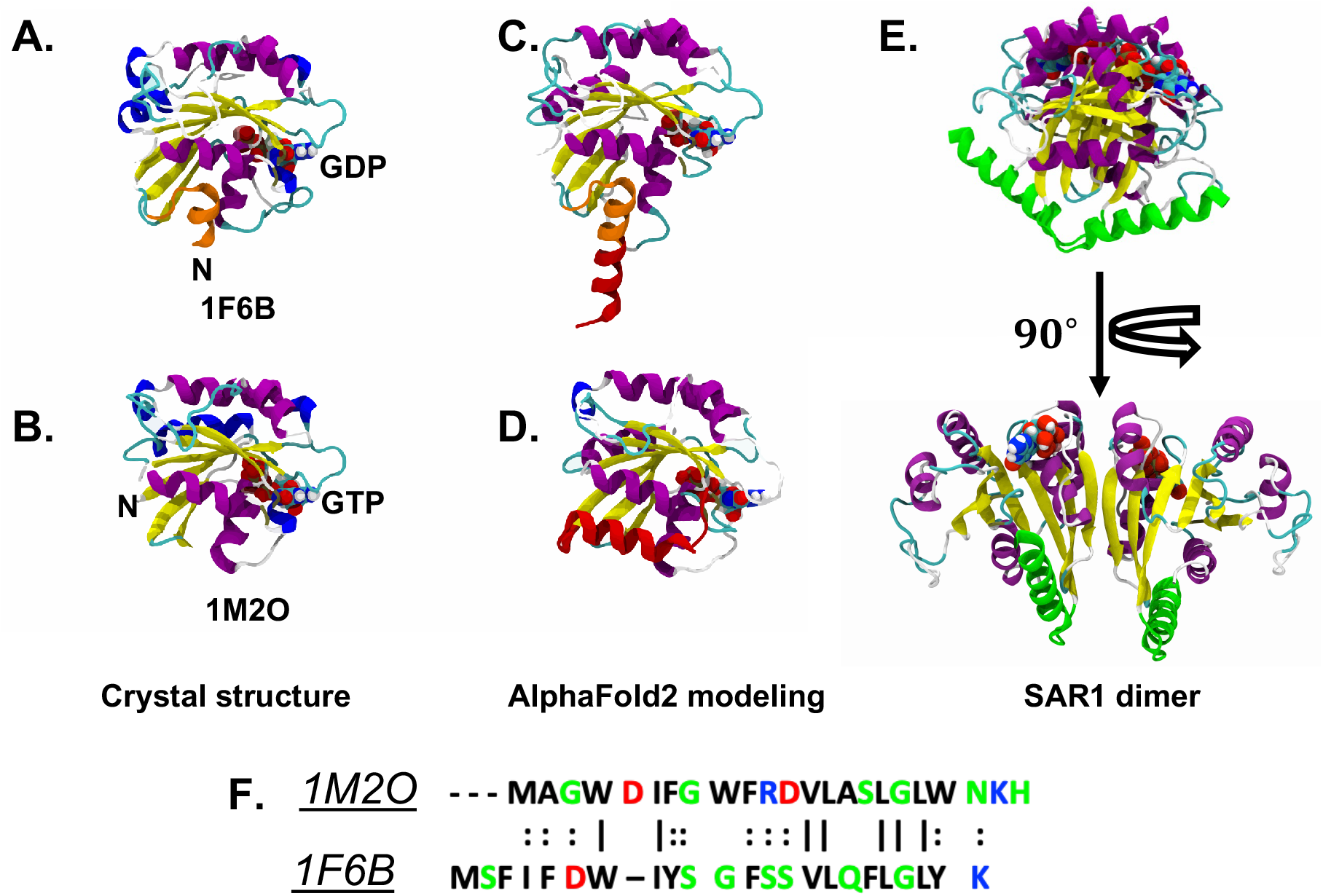
Structural features and sequence of Sar1. (A-B) Crystal structures of Sar1 in the GDP-(PDB id: 1F6B) and GTP-(PDB id: 1M2O) bound states shown as cartoon representations. GDP and GTP are shown in VDW. Residues 1-12 in the GDP-bound state are highlighted with orange color. (C-D) h-GDP (top) and y-GTP (bottom) models of Sar1 derived from AlphaFold2. The amino-terminal region, which is missing in the original crystal structures, are predicted to be helical and shown in red. (E) A dimer model of Sar1 obtained from the h-GTP state with the helical amino-terminus shown in green. (F) Aligned sequence of Sar1 in y-GTP (1M2O) and h-GDP (1F6B) states with the following color codes, acidic: red; basic: blue; polar: green; non-polar: black. Vertical line represents sequence identity while double dots indicate sequence similarity.

In contrast, with the first 12 residues in the Sar1-GDP structure and the first 23 residues in the GNP (GTP)-bound structure remaining unresolved, it remains unclear whether the noted structural differences in the amino-terminal region are sufficient to explain the membrane sculpting ability of Sar1 in its different nucleotide-bound states. For example, our previous work (14) examining the effect of nucleotide binding on Sar1 membrane association indicated that, in solution, the amino-terminus was not buried inside the protein core in the GDP-bound state as hypothesized based on the crystal structure(9). Further, we observed that GDP-bound Sar1 remains capable of membrane binding, although its affinity is not as strong as that for the GTP-bound state. Evidently, the structural impact of nucleotide binding on Sar1 and its contributions to membrane remodeling requires a better molecular understanding.

Atomistic and coarse-grained Molecular Dynamics (MD) simulations have played a pivotal role in understanding the molecular mechanism of protein-induced membrane curvature generation.(15) For example, the BAR domain is one of the most well studied proteins in this context and has been shown to induce membrane curvature through both scaffolding and amphipathic helix insertion(16–20). Recently, atomistic and coarse-grained MD simulations were employed to elucidate the molecular mechanism of membrane invagination and cell division driven by the Endosomal Sorting Complex Required for Transport III (ESCRT-III).(21–23) Further, combined all-atom and continuum elastic model-based calculations indicated that spontaneous curvature of an inclusion varies non-monotonically with the inclusion height of the bound protein/peptide.(24) Nevertheless, to date, an understanding of how amino acid sequence and penetration of a membrane binding protein motif controls the membrane curvature generation process remains incomplete. This is in part due to limitations in the scale of conformational variations feasible in the membrane binding region of proteins, creating challenges to characterizing differences in their membrane bending activities. In this context, Sar1 serves as a suitable system for probing the effect of relatively subtle variations in the amino-terminal structure and sequence on its distinct abilities to penetrate and sculpt lipid membranes in different nucleotide-bound states, which ultimately regulate COPII-mediated protein trafficking.

Specifically in this study, we employ atomistic MD simulations to provide a comparative assessment of membrane insertion and curvature generation by Sar1 in the GDP- and GTP-bound states. We also investigate how dimerization of Sar1 in the GTP-bound state affects its membrane remodeling activities. First, we model the missing coordinates of the amino-terminal region of Sar1 using AlphaFold2(25) and perform solution simulations to characterize its dynamics in different states (GTP vs. GDP vs. dimer) in the absence of membrane. Then, we quantify the membrane binding ability of Sar1 in terms of penetration depth and number of contacts. Finally, using a membrane ribbon protocol(21, 26), we assess membrane bending ability of Sar1 in different states. The results support a molecular mechanism in which GTP binding induces stable penetration of Sar1 into the membrane through its complete amino-terminal helix, leading to an intense degree of membrane bending. Dimerization of the GTP-bound state further enhances the amount of positive curvature induced on the membrane, since both amino-terminal helices contribute. In the GDP-bound state, only a small portion of the amino-terminus is available for interactions with the membrane, which results in weaker membrane binding(14) and less membrane curvature generation. Overall, by comparing multiple simulations, we provide an explicit correlation between the degree of membrane bending and the amount of protein volume inserted into the membrane, whereas the membrane penetration depth alone is unable to explain the trend of protein-induced membrane curvature.

## Results

### A. GTP binding increases the flexibility of the amino-terminal region of Sar1

Before examining its interactions with membrane, we first assess the dynamics of Sar1 in solution, focusing on the amino-terminal segment of Sar1 that acts as the membrane anchor. For this purpose, we employ an energy landscape sensitive metric known as Cumulative Variance of Coordinate Fluctuations (*σ*^2^_*CVCF*_), which was previously shown to characterize the variation in dynamics even when average structural changes are limited.(27) It reports the curvature of the underlying energy landscape and is able to detect local convergence of sampling when Boltzmann population is achieved in the sampled energy minima. For the analysis, we choose the set of backbone atoms of the amino-terminus, which is divided into two segments: residues 1-12 and residues 13-23; the latter was observed to retract into a surface pocket formed partly by the loop connecting the *β*2-*β*3 hairpin in the GDP-bound state and believed to be solvent-exposed in the GTP-bound state.

Our analysis reveals that backbone *σ*^2^_*CVCF*_ is significantly higher in the y-GTP state compared to the h-GDP state for both segments (Fig-2). For example, backbone *σ*^2^_*CVCF*_ of residues 1-12 is 255 Å^2^ for the h-GDP state and rises to 950 Å^2^ in the y-GTP state (Fig-2A top). Similarly, for residues 13-23, backbone *σ*^2^_*CVCF*_ increases from 35 Å^2^ to 91 Å^2^ (Fig-2A bottom). In both nucleotide-bound states, residues 1-12 exhibit increased flexibility (by a factor ~10 in *σ*^2^_*CVCF*_) than residues 13-23. Similar trends in flexibility are observed with the CHARMM-GUI derived structural models (Fig-2B); the absolute values of *σ*^2^_*CVCF*_ are substantially higher (Fig-2B) since the amino-terminal region in the CHARMM-GUI models is disordered (Fig-S1) rather than being helical as in the AlphaFold2 models (Fig-1C-D). In the homology model for the hamster GTP-bound state, h-GTP, both segments exhibit higher *σ*^2^_*CVCF*_ values than the h-GDP-bound state. In the h-GTP-bound dimer, the amino-terminal helices generally become less flexible, especially residues 13-23; nevertheless, they exhibit higher fluctuations than the monomeric GDP-bound state (Fig-2C).

**Fig. 2.**
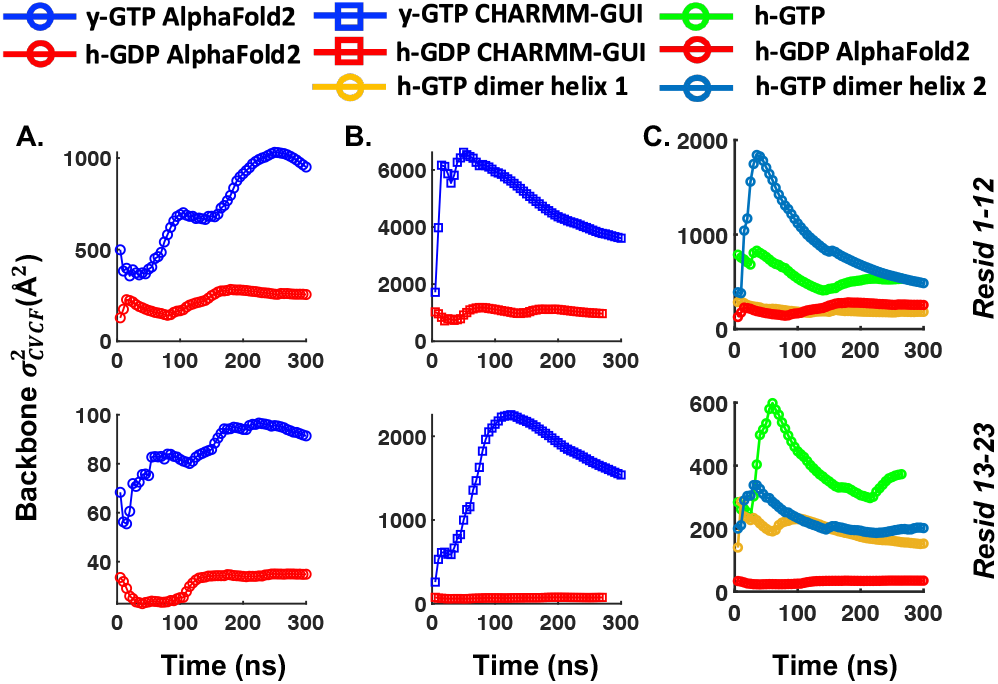
Assessment of the amino-terminal flexibility of GDP- and GTP-bound Sar1 in different models in solution. (A) Backbone CVCF traces of the amino-terminal residues 1 to 12 (top) and 13 to 23 (bottom) of the AlphaFold2 models. (B) Similar plot for Sar1 where the amino-terminus is modelled with CHARMM-GUI. (C) Comparison of amino-terminal flexibility of the h-GTP Sar1 model with the h-GDP model from AlphaFold2. Backbone CVCF traces of each of the helices of the h-GTP dimer model are also shown for comparison.

To better understand how GTP binding induces enhancement in the flexibility of the amino-terminal segment, we computed C*α*-C*α* contacts (10 Å cutoff) averaged over trajectory frames. GDP-bound structures form excess contacts near the amino-terminal region that are lacking in the GTP-bound cases (compare Fig-3A vs. Fig-3B, Fig-3D vs. Fig-3E). These differences reflect the compact nature of the amino-terminus in the GDP-bound state, while in the GTP-bound state, the amino-terminal segment remains largely detached from the protein core. As a result, the amino-terminus forms more contacts with the carboxyl-terminus in the GTP-bound state in solution (black arrow in Fig-3B,C). In the case of the h-GTP-bound dimer, residues 13-23 of helix 1 form stronger contacts with residues 65-70 compared to the monomeric state; in addition, the first segment (residues 1-12) of helix 2 rapidly switches from an open to a closed state and forms extensive contacts with residues 159-163 (Fig-3C vs. Fig-3F), as also reflected by the large rise in *σ*^2^_*CVCF*_ followed by a sharp decay (Fig-2C top). Taking the CVCF values and the contact maps together, regardless of the structural models, the amino-terminus in the GTP-bound state exhibits a larger conformational heterogeneity that may facilitate the process of membrane binding and penetration.

**Fig. 3.**
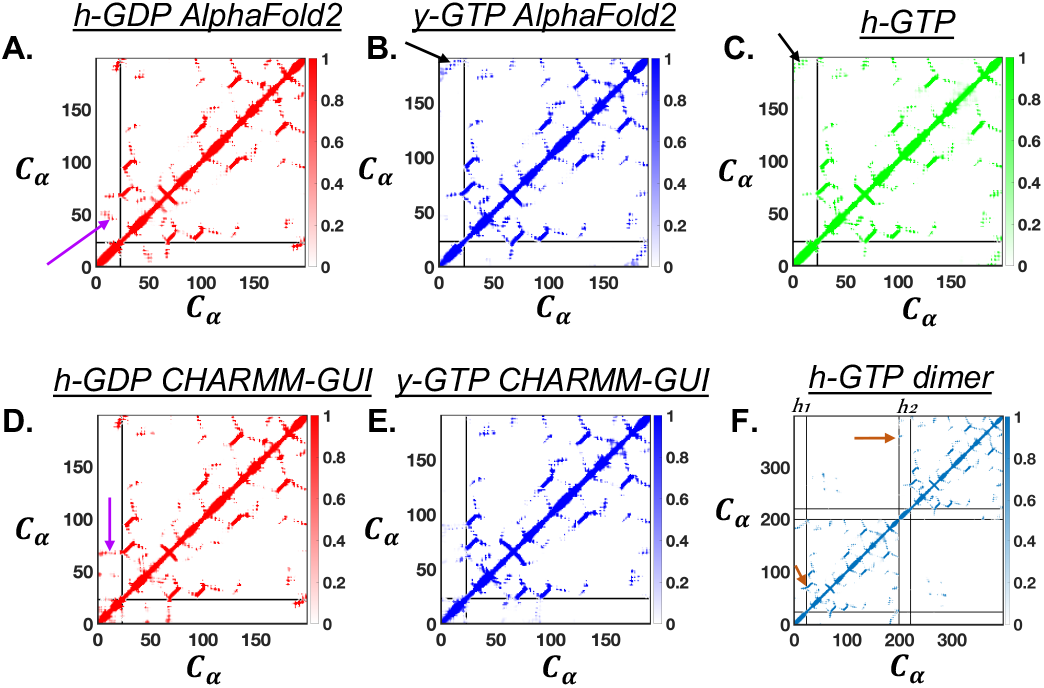
Trajectory averaged C*α*-C*α* contact analysis (10 Å cutoff) of AlphaFold2 derived (A) h-GDP (B) y-GTP and (C) h-GTP models of Sar1. Similar analysis for the CHARMM-GUI derived (C) h-GDP (D) y-GTP models of Sar1. (E) Contact map of the h-GTP dimer considering all C*α* atoms. The black lines in A-E indicate the position of residue 23, which is the last residue in the amino-terminal motif. In addition to the residue 23 of helix 1, residues 1 and 23 of helix 2 of the h-GTP-bound dimer are also marked by black lines in (F). The purple arrow indicates the excess contacts in the GDP-bound structures which arise due to the more compact amino-terminus compared to the GTP-bound structures. The extended amino-terminus of GTP-bound structures often makes contacts with the carboxyl-terminus as indicated by the black arrow. Brown arrow indicates the excess contact made by the amino-terminal helices in the h-GTP-bound dimer that are not present in the corresponding monomer state.

### B. Sar1 binds tighter to and penetrates deeper into the membrane in the GTP-bound state

Sar1 readily binds to the membrane with the HMMM model at a higher (1.7 times) area per lipid. Despite being placed at ~ 5 – 15 Å away from the membrane, Sar1 inserts into the membrane in almost all (9 out of 10) cases. Since Sar1 has multiple hydrophobic residues (13 in y-GTP and 14 in h-GDP) in its amino-terminus, a significant membrane binding affinity is to be expected.

However, almost all amino-terminal residues penetrate into the membrane in both h-GDP- and y-GTP-bound states (Fig-S3), which exhibit limited difference in membrane binding in HMMM simulations at expanded area per lipid values. The only notable difference is that the unstructured amino-terminus in the CHARMM-GUI models acquires helicity upon insertion into the membrane in the y-GTP-bound state (Fig-S4 A) but not in the h-GDP-bound state (Fig-S4 B), which is probably due to the longer contiguous helix in the former system.

When the HMMM model is converted into full lipid membrane, differences in Sar1-membrane interactions in different nucleotide bound states become more prominent. In the following, we discuss several representative conformations of the AlphaFold2 h-GDP and y-GTP models for their detailed interactions with the full membrane model.

While the precise depth of membrane penetration varies for each nucleotide-bound state with simulations on the order of 80-160 ns, the mass density profiles suggest that, overall, Sar1 penetrates deeper into the membrane when bound to GTP. With the h-GDP model, residues 13-23 do not enter the membrane as indicated by the broad mass density distributions (see brown arrow in Fig-4B-C). By contrast, y-GTP-bound Sar1 leads to complete insertion of the amino-terminal anchor into the membrane, leading to much narrower mass density distributions (black line in Fig-4D-E). For example, with conformation 1 of the h-GDP-bound state, the mass density peaks (black dotted line in Fig-4B) coincide with that of the C2 atom of DOPC. With the deepest insertion of the y-GTP-bound state, the mass density of the amino-terminus fully covers that of the C24 lipid atom (yellow curve in Fig-4D) and its decay coincides with that of the C27 lipid atom (magenta curve in Fig-4D), which is close to the membrane core (cyan dotted line). The latter observation leads to an estimated insertion depth (*d*) of ~ 1.12 *nm* (see Methods and sec. 1E in SI text for details) for the GTP-bound state, which is comparable to the value of 1-1.5 *nm* obtained in fluorescence quenching experiments by Hanna et al.(14).

**Fig. 4.**
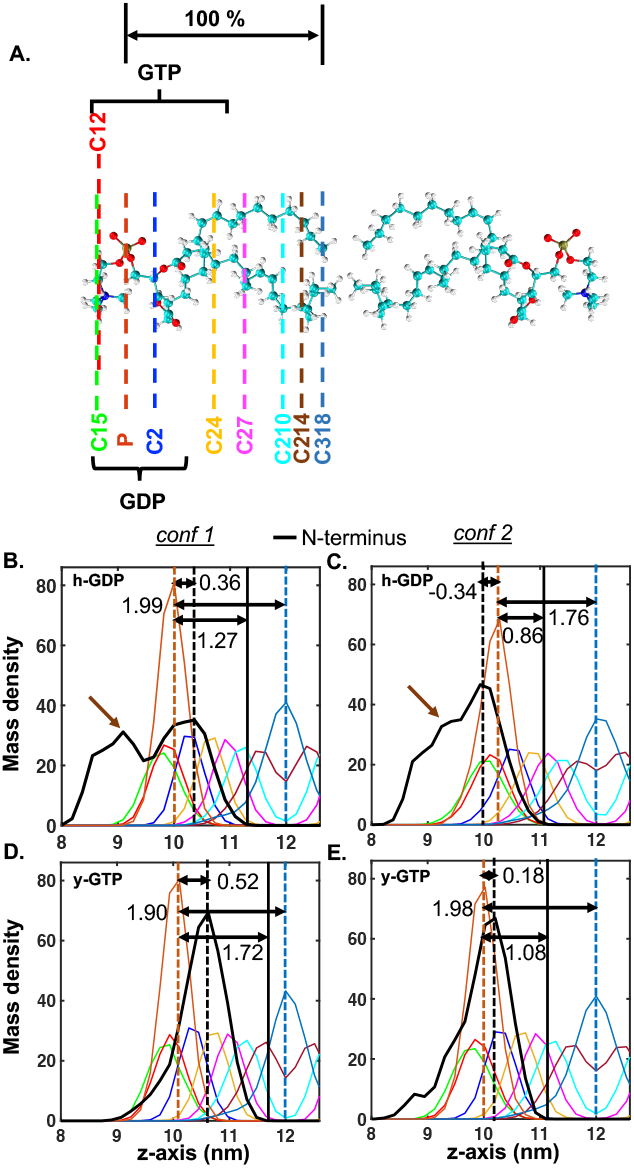
Assessment of membrane penetration depth of AlphaFold2 modelled Sar1 as a function of bound nucleotide state. (A) A pair of DOPC molecules from upper (left) and lower (right) leaflet of the membrane. Mass density profile of various atoms of the DOPC lipid molecules (shown in A) and the amino-terminal region (residues 1-23) of Sar1 in the presence of (B-C) GDP and (D-E) GTP seeded from two different initial conformations. Mass densities of individual lipid atoms are color coded (see A) and the protein amino-terminus is shown in black. The phosphate plane is considered as the membrane periphery and is highlighted by a brown dotted line. The black dotted line represents the peak position of the amino-terminal anchor and the cyan dotted line indicates the core of the membrane. The black solid line marks the region inside membrane where mass density of amino-terminal anchor decays to 0. For the h-GDP-bound state, a significant portion of the amino-terminus remains outside of the membrane as pointed out by the brown arrow. We normalize the penetration depth with the membrane leaflet thickness defined by the distance between membrane core (C318) and periphery (phosphate).

We provide snapshots of membrane-bound Sar1 in different nucleotide states to illustrate differential insertion of the amino-terminus into the membrane (Fig-5A-B). Residues 13-23 remain unbound to the membrane for h-GDP-bound SAR1 as shown in Fig-5A (few residues represented in VDW away from the membrane). In contrast, the full amino-terminus remains inserted into the membrane for y-GTP-bound SAR1 (no residues represented in VDW away from membrane in Fig-5B). Partitioning of polar (green, red and blue) and non-polar (black) residues at the membrane/water interface is observed in both cases. In the h-GDP-bound state, Met1, Phe3, Ile4, Phe5, Trp7 and Ile8 remain buried inside the membrane, and Ser2, despite being polar, also stays inside the membrane. Tyr9 and Phe12 along with the rest of the polar residues among residues 1-12 remain solvent exposed. In y-GTP-bound Sar1, all hydrophobic residues of the amino-terminus remain inserted where Ile6 penetrates the membrane most deeply; polar, acidic and basic residues stay at the surface of the membrane, pointing outwards from the membrane.

**Fig. 5.**
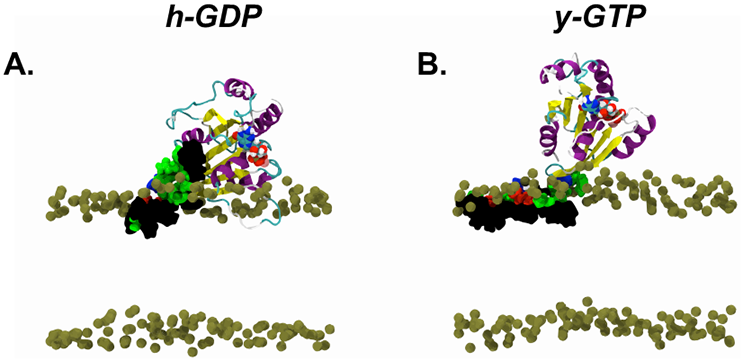
Snapshots of the conformation 1 of Sar1 in (A) h-GDP- and (B) y-GTP-bound states absorbed onto the fully atomistic membrane after 160 ns of simulation. Amino-terminal residues (residues 1-23) of Sar1 are shown in VDW with the similar color code as shown in Fig-1F. GTP and GDP are shown in the van der Waals representation. Only phosphate atoms of the membrane are shown for clarity.

To further quantify the contrasting membrane binding activities for different nucleotide-bound states of Sar1, we computed the total number of contacts between the membrane and the protein amino-terminal segments. While residues 1-12 form close contacts with the membrane in both nucleotide-bound states (Fig-6A), differences arise in the contacts between residues 13-23 and the membrane; for example, the y-GTP state exhibits significantly higher numbers of contacts (~ 1500 in conf 1 and ~ 2000 in conf 2) with the membrane through residues 13-23 (Fig-6B blue) compared to the h-GDP case (< 500, as shown in Fig-6B red).

**Fig. 6.**
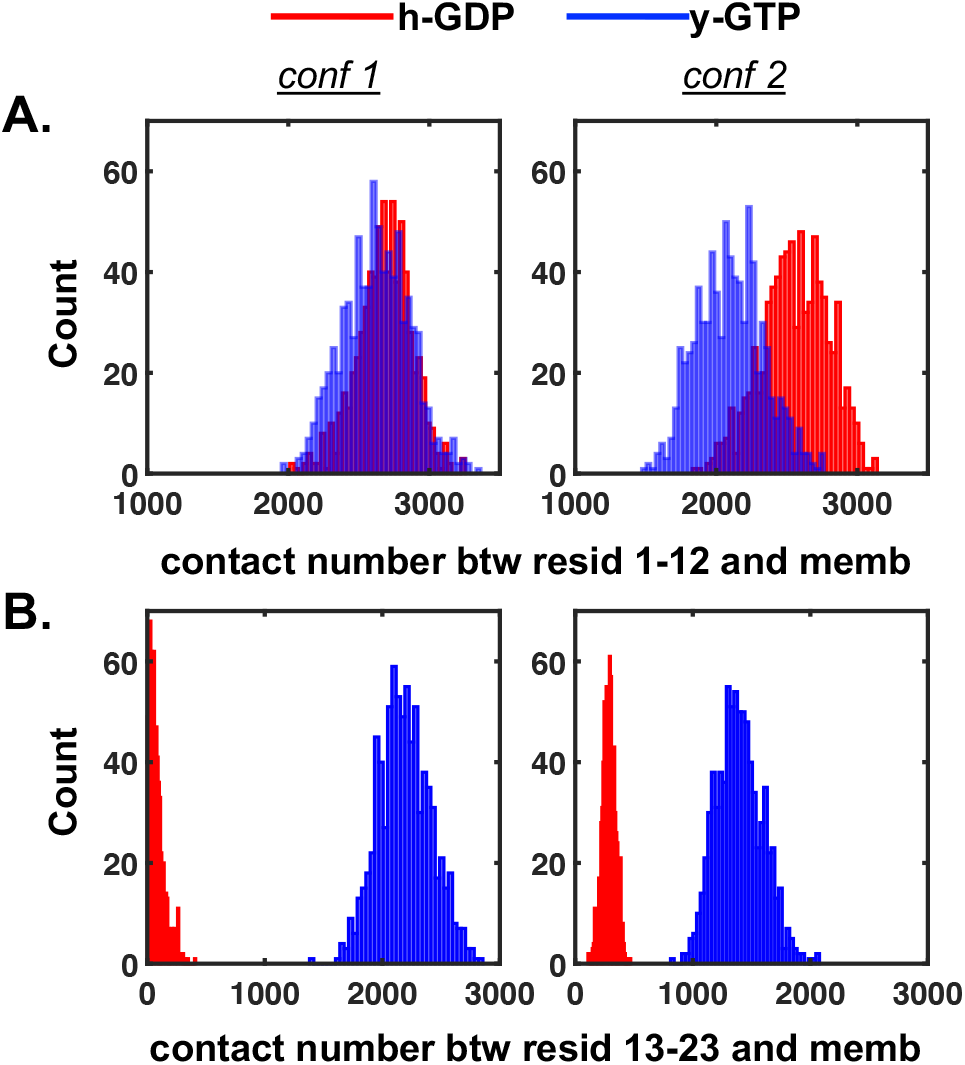
Protein-membrane contact analysis. Histograms of the total numbers of contacts (defined based on atomic distances less than 5 Å) between membrane and the amino-terminal residues of Sar1 in GDP-(red) and GTP-(blue) bound states obtained from 80-160 ns of simulations; for each nucleotide state, results for two conformations (as shown in Fig. 4) are shown (conformation 1 in left and conformation 2 in right). The amino-terminus is divided into two parts: residues 1-12 (A) and 13-23 (B).

### C. Sequence dissimilarities between h-GDP- and y-GT-P-bound Sar1 do not contribute to the differential membrane activities

As indicated in the Introduction, crystal structures for the GDP- and GTP-bound forms of Sar1 were derived from different organisms; the sequence identity and similarity between them is 56 % and 69 %, respectively. The amino-terminal segments, which act as the membrane anchor, also bear sequence differences (Fig-1E). The y-GTP structure contains 2 acidic and 2 basic residues, while the h-GDP structure has 1 acidic and 1 basic residue. Nevertheless, the amino-termini remain amphipathic in both cases.

To demonstrate that the origin of the contrasting membrane binding activities of h-GDP- and y-GTP-bound Sar1 lies primarily in the distinct structures of the amino-terminus rather than in the sequence dissimilarities, we built the homology model h-GTP structure (see Methods), which takes the sequence of h-GDP-bound state but uses the y-GTP-bound state as the structural template. The h-GTP structure exhibits complete insertion of the amino-terminus into the membrane (Fig-S6 A), thus leading to the burial of Val15, Leu16, Phe18, Leu19, Leu21 and Tyr22, in addition to the hydrophobic residues from residues 1-12. The mass density profile (Fig-S6 B) indicates that the amino-terminus penetrates beyond the C2 atom of DOPC, similar to the case of conformation 1 of y-GTP (Fig-S6 C); the numbers of membrane contacts also indicate strong membrane binding of the h-GTP-bound model for both residues 1-12 and 13-21 (Fig-S6 C-D). Another conformation of h-GTP yields slightly less penetration depth (Fig-S7 A) but a similar number of membrane contacts (Fig-S7 B-C). Therefore, the h-GTP-bound state resembles y-GTP in terms of the membrane binding and penetration activities, confirming that the structural features of the amino-terminal helix are more important than the precise sequence.

### D. Dimerization of Sar1 impacts the protein membrane interactions

In EM studies, Sar1 was observed to form lattice assemblies on constricted membrane surfaces where the smallest repeating unit was a dimer.(28) Hence, it is important to understand the impact of Sar1 dimerization on its interaction with the membrane. In the simulation, both amino-terminal helices exhibit less penetration depth (0.81, 0.73 nm) into the membrane (Fig-7B-C) compared to the monomeric h-GTP model (Fig-S6 B); this is likely due to the reduced flexibility of amino-terminal helices in the dimer (Fig-2C). Correspondingly, each of the amino-terminal membrane anchors forms a smaller number of contacts with the membrane compared to the monomeric case (Fig-7 D-E). However, considering contributions from both helices, the Sar1 dimer forms a significantly higher number of contacts (~ 8000 vs. 5000) with the membrane as compared to a monomer. Overall, dimerization of Sar1 significantly impacts its interactions with the membrane, which in turn influences its membrane bending ability (*vide infra*).

**Fig. 7.**
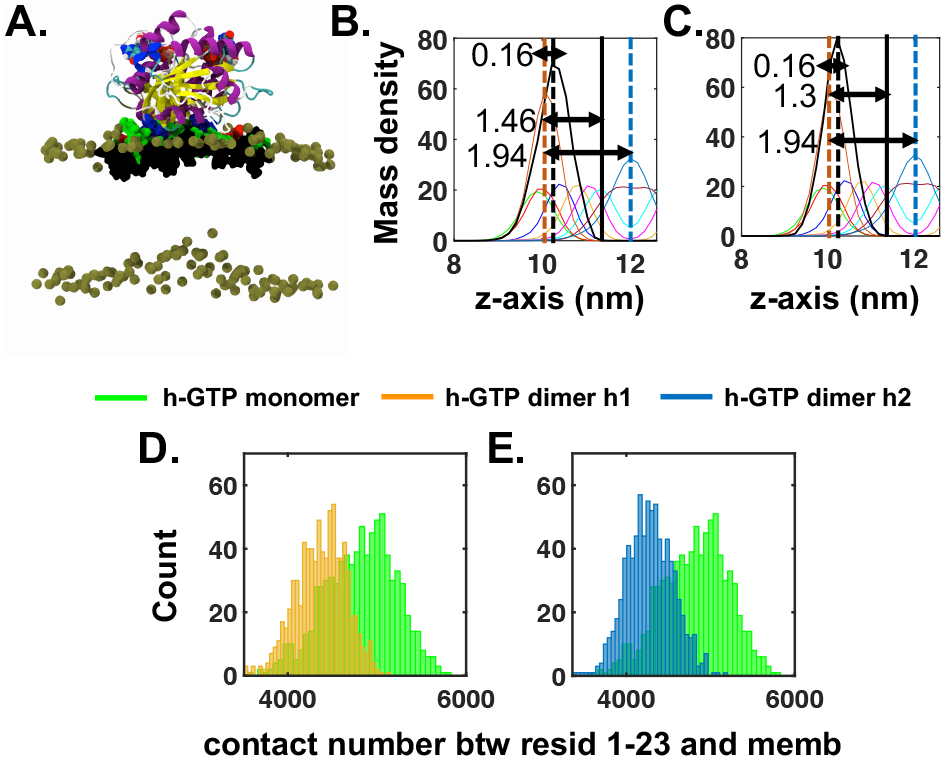
Membrane penetration depth and the number of contacts with membrane in Sar1 dimer. (A) Snapshot of h-GTP-bound dimer model of Sar1 embedded in the membrane after 160 ns of simulation. Residues 1-23 of the both the helices are shown in VDW representation with the color code similar to that in Fig-1F and Fig-5. Mass density profile of various atoms (color coded as shown in Fig-4A) of DOPC molecules along with that (in Black) of (B) amino-terminal helix 1 and (C) amino-terminal helix 2 of a Sar1 dimer. (D-E) Histograms of the total numbers of contacts (definition similar to that in Fig-6) between membrane and the amino-terminal residues of Sar1. The amino-terminal helices of dimers individually form fewer contacts than that of monomers (shown in green) but their total number of contacts far exceeds that of monomer.

### E. GTP binding and dimerization significantly enhance the ability of Sar1 to generate positive curvature on membrane

To assess the impact of the nucleotide-bound state and dimerization on the membrane sculpting ability of Sar1, we perform membrane ribbon simulations. For each case, we start with the conformation that exhibits the deepest membrane insertion (as discussed in the previous subsection) on a large membrane patch of ~ 32 nm length and conduct curvature generation simulations (see Methods, sec. 1F in SI text and Fig-S8).

During the simulation, the membrane ribbon elongates along the *X* axis and gradually adopts a positively curved shape (Fig-8A-F) within ~ 15-20 ns in case of the conformation 2 of h-GTP-bound Sar1; the membrane midplane reaches up to ~ 2 nm shifts (magenta line in Fig-8G), which correspond to a mean curvature of 0.016 nm^-1^(Fig-S10-S12). The conformation 1 of y-GTP model leads to slightly higher membrane bending where the membrane midplane goes beyond ~ 2 nm (blue line in Fig-8G), which corresponds to a positive membrane curvature of 0.019 nm^-1^. Further, conformation 1 of h-GTP-bound Sar1 yields an even higher degree of positive curvature (0.023 nm^-1^, green line in Fig-8G), while conformation 2 of y-GTP-bound Sar1 induces a moderate positive curvature (0.0046 nm^-1^, brown line in Fig-8G). By contrast, h-GDP-bound Sar1 does not lead to any significant impact (curvature ~ 0.0011 nm^-1^) on the membrane morphology (red line in Fig-8G); we did not pursue membrane ribbon simulation with conformation 2 of h-GDP model since it yields poor penetration into the membrane (mass density peak of the protein amino-terminus lies outside the phosphate plane in Fig-4B). As discussed above, conformation 1 of y-GTP and conformation 1 and 2 of h-GTP-bound forms of Sar1 insert into the membrane to a greater extent than the h-GDP-bound state (Fig-4, 5, 6, S6 and S7). These differences in the penetration depth arise due to the insertion of a higher number of hydrophobic residues into the membrane in the GTP-bound state (8 in GDP- vs. 13 in GTP-bound Sar1). The membrane ribbon simulations in turn reveal that such enhancement in penetration depth and the number of hydrophobic residues inside the membrane leads to significant amplification (~ 10-20 times) in positive membrane curvature generation. We note that there are 13 hydrophobic residues in the y-GTP-bound Sar1 and 14 in the h-GTP-bound Sar1; the larger number of hydrophobic residues in the latter case may contribute to its generation of slightly enhanced (blue vs. green line in Fig-8G) membrane curvature. Further, the h-GTP dimer model of Sar1 induces an even stronger positive curvature where the membrane midplane reaches ~ 5 nm shifts (azure line in Fig-8G), which corresponds to a mean curvature of 0.0309 nm^-1^. Evidently, despite exhibiting shallower penetration depths (Fig-9B-C), the Sar1 dimer produces an intense bending of the membrane much stronger than a monomer.

**Fig. 8.**
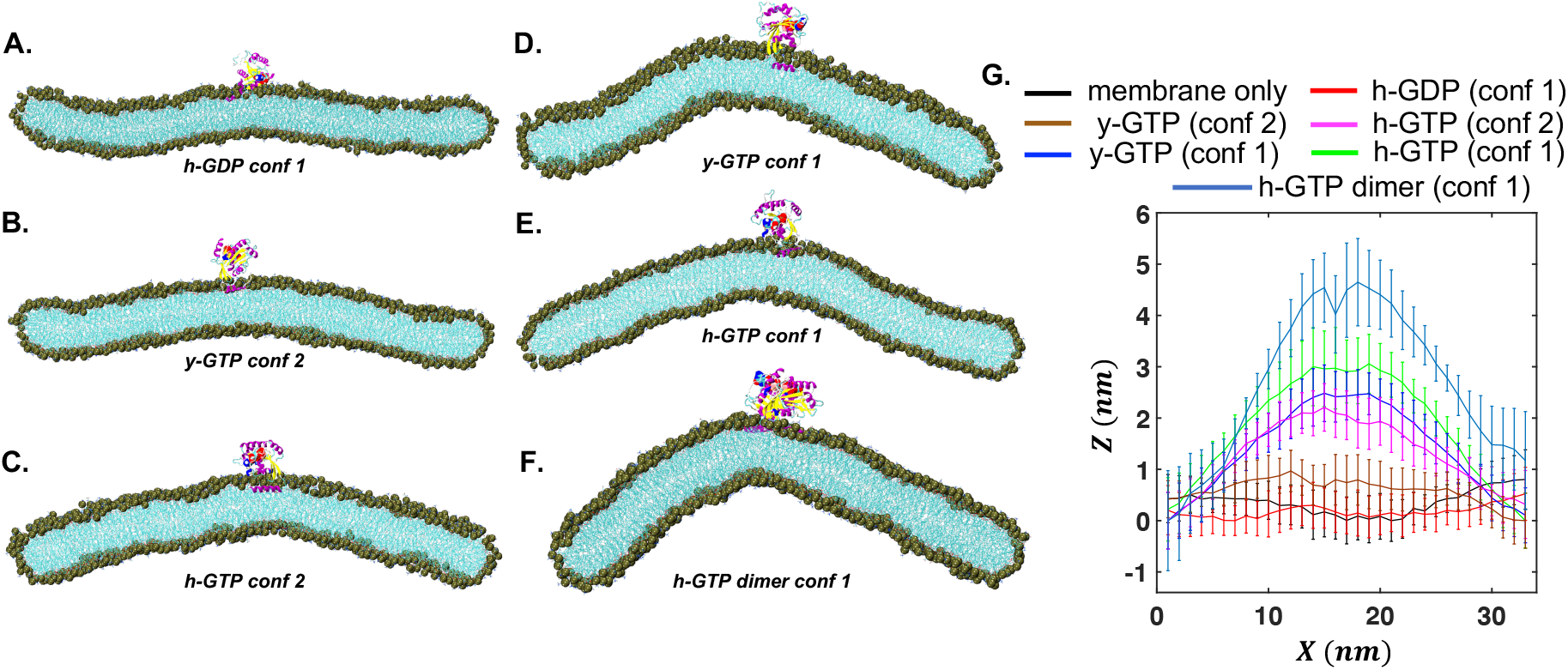
Membrane ribbon simulations of different models of Sar1 in the GDP- and GTP-bound states. Snapshots of (A) conformation 1 of h-GDP, (B) conformation 2 of y-GTP, (C) conformation 2 of h-GTP, (D) conformation 1 of y-GTP, (E) conformation 1 of h-GTP, and (F) conformation 1 of h-GTP dimer model of Sar1 in the presence of membrane. (G) The *Z*-positions of the membrane mid-plane averaged over *Y*-direction as a function of the *X* position for the various membrane ribbon simulations. While the h-GDP-bound state does not produce any significant mean curvature, all GTP-bound states (h-GTP and y-GTP models) lead to significant positive mean curvature in the membrane ribbon along *X*. This positive curvature is further enhanced with the binding of a Sar1 dimer.

**Fig. 9.**
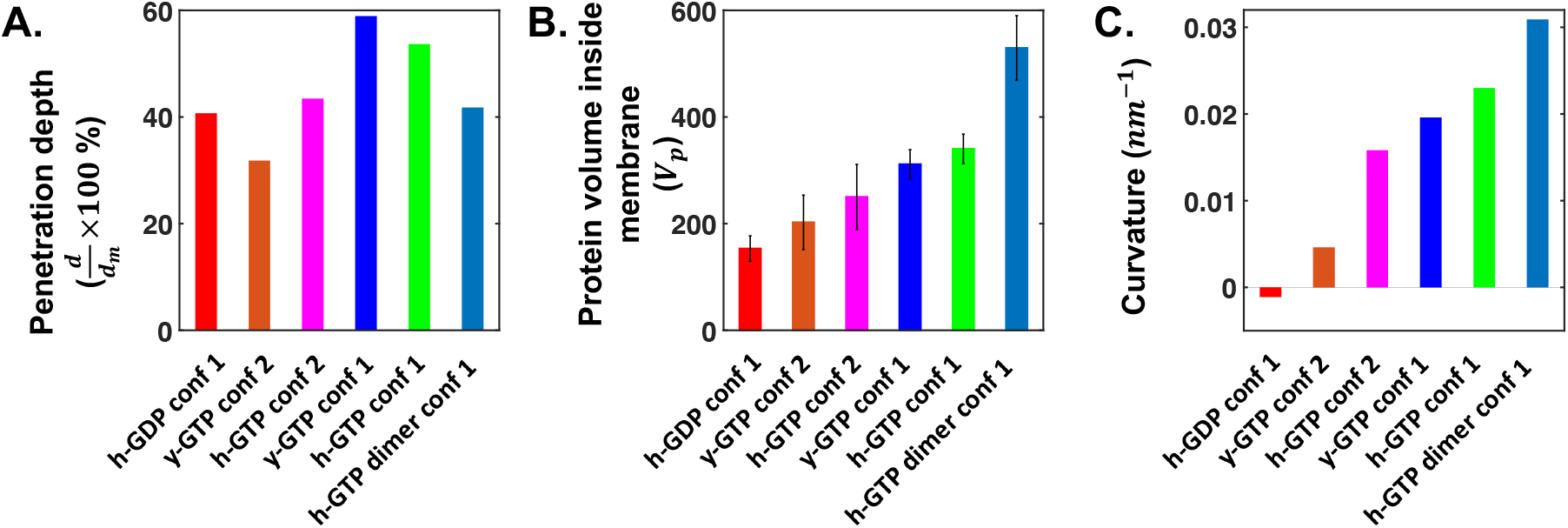
Correlation between membrane curvature generation and insertion volume of protein. Bar plot of (A) membrane penetration depth (B) inserted protein volume (*V_p_*) in the membrane and (C) the magnitude of curvature induced on the membrane for various models of Sar1. The extent of curvature generation correlates well with the number of protein atoms inserted into the membrane while membrane penetration depths of the proteins do not follow the trend well.

To better understand the factors that determine the degree of membrane bending, we compare the penetration depth and protein volume inserted into the membrane with the induced curvature. As shown in Fig. 9, the degree of generated membrane curvature is not solely dependent on the penetration depth of the amphipathic helix; rather, the curvature is well correlated with the number of protein atoms inserted into the membrane, which reflects the volume (*V_p_*) of protein insertion. For example, conformation 2 of h-GTP exhibits a similar membrane insertion depth as conformation 1 of h-GDP (Fig. 9A); however, they differ significantly in the number of protein atoms inserted into the membrane (Fig. 9B), since the full amino-terminal helix is involved in the h-GTP-bound state, in contrast to the first 12 residues in the h-GDP-bound state. The latter trend is better correlated with the significantly higher membrane curvature generated by the h-GTP-bound Sar1 model (Fig. 9C). Moreover, the penetration depth of Sar1 dimer is notably less than that of the monomer but the two amino-terminal helices of the dimer causes much higher protein volume inclusion inside the membrane, leading to a significant membrane curvature (however, for a relevant technical point of the membrane ribbon protocol, see Fig. S9).

## Discussions

In this study, we provide an atomistic description of the Sar1/membrane interface and characterize its modulation as a function of the bound nucleotide state. Previous *in vitro* and *in vivo* studies(4, 29, 30) showed that the amino-terminal region of Sar1 acts as an essential factor in the generation of COPII-coated transport carriers. Further, Sar1 performs its function in a GTP dependent manner, in which GTP binding activates Sar1 to begin the membrane bending process, while GTP hydrolysis facilitates detachment of carriers from the ER membrane. The crystal structures of Sar1 suggested that GTP binding triggered the exposure of the amino-terminus to facilitate its membrane bending activity.(9) A detailed comparison of the two crystal structures reveals that the *β*2-*β*3 hairpin is displaced by ~ 7 Å as a consequence of GTP binding, resulting in alteration of hydrogen bonding between *β*1-*β*3. Consequently, the surface pocket that holds the amino-terminal membrane anchor in the GDP-bound state is eliminated, which releases the amino-terminus to be available for membrane binding. Recent Cryo-EM experiments in combination with Rosetta based modeling also indicated a parallel insertion of the entire amino-terminal amphipathic helix in the GTP bound state of SAR1.(31) However, fluorescence quenching-based *in vitro* measurements indicated that at least part of the amino-terminal anchor remained solvent exposed irrespective of the bound nucleotide state.(14) In fact, both GTP and GDP bound forms of Sar1 exhibited significant reduction in fluorescence quenching in the presence of membrane, implying membrane association of Sar1 in a nucleotide independent fashion. Our simulation results support that Sar1 binds to the membrane in both GDP- and GTP-bound states (Fig-4–5). At the quantitative level, the extent of reduction in fluorescence quenching in the presence of membrane was different in GDP- and GTP-bound forms of Sar1; GTP-bound Sar1 showed significantly stronger binding to the membrane compared to the GDP-bound state, which caused the lowering of the Stern-Volmer constant value by ~ 2.7 times.

Our simulations provide an atomistic explanation of the experimental observations by showing that GTP-bound Sar1 penetrates well into the membrane by anchoring its full amino-terminus (residues 1-23), while the latter only partially inserts into the membrane in the GDP-bound state (Fig-4B vs. 4D). Indeed, contact analysis reveals that residues 13-23 remain outside of the membrane in the GDP-bound state, while they are immersed in the membrane in the GTP-bound state (Fig-6A-B). Moreover, we find penetration of GTP-bound Sar1 up to C27 atoms of the lipids, which lie just beneath the hydrophilic region of the membrane; this corresponds to an insertion depth of ~ 1.12-1.18 nm, which compares favorably with the experimental estimate of 1-1.5 nm(14).

The membrane ribbon simulations capture ~ 10-20 times enhancement in the membrane curvature generation (Fig-8) as a result of relatively subtle changes in the conformation of the amino-terminal membrane anchor of Sar1 due to change in the nucleotide-bound state. This increased ability of membrane bending originates from the more extensive membrane penetration of the amino-terminus in the GTP-bound state. The membrane ribbon simulation (Fig-8 and S6) shows that a single GTP-bound Sar1 is able to produce 42.71-51.13 *nm* radius of curvature on the membrane; this value is comparable to that generated by N-BAR domains (radius ~ 54 *nm*) and Vps32 protomers of the ESCRT-III complex (radius 30.5 ± 9.5 nm).(21) The observed curvature is also consistent with experimental studies, which reported the formation of COPII-coated carriers of 25-50 *nm* radius.(32, 33) Sar1 has been shown to assemble on membranes and cause membrane constriction following tubule formation.(28, 29). The membrane associated Sar1 lattice structure reveals that multiple dimer units of Sar1 co-assemble and form ordered complexes. Another experimental study also suggests that the modulation of membrane mechanical properties is proportional to the concentration of Sar1 dimers(34). Thus, the dimeric state of Sar1 bear immense functional importance(8). Our study explicitly demonstrates that the Sar1 dimer is able to generate much stronger membrane bending compared to the monomeric state. In polymeric Sar1 assemblies, it is possible that the induced membrane curvature is further enhanced, although the effect might be sensitive to the relative orientations of the Sar1 dimers and therefore needs to be analyzed explicitly in future studies.

Our explicit comparison of various states of Sar1 in terms of their membrane bending activities provides insights into factors that directly impact the degree of membrane curvature generation. For example, while the correlation between insertion depth of membrane inclusion and curvature generation has been shown with continuum mechanics models(35), we note that the degree of membrane bending does not depend monotonically on the insertion depth. As shown in our recent coarse-grained analyses, deep insertion of protein motifs may lead to negative rather than positive membrane curvature(22). In this study, we estimate membrane penetration depth for each of the 6 Sar1 models (Fig-9A). An interesting observation is that the trend in membrane penetration depth does not match with that in the degree of membrane bending (Fig-9C). Instead, the magnitude of membrane curvature generation correlates well with the volume of protein (*V_p_*) inserted inside membrane (Fig-9B vs. Fig-9C). These observations establish protein volume inserted into the membrane as the key predictor for membrane curvature generation.

In classic models for COPII-mediated trafficking, Sar1 recruits Sec23-Sec24 to form a pre-budding complex, and the resulting Sar1-Sec23-Sec24 inner-coat complex in turn recruits Sec13-Sec31 to produce the outer-coat layer. Finally, GTP hydrolysis on Sar1 triggers the detachment of transport carriers from the ER membrane. Based on our penetration depth analysis of GDP- and GTP-bound states of Sar1, we anticipate that upon GTP hydrolysis, the amino-terminal membrane anchor becomes retracted from the membrane, which, as shown explicitly by membrane ribbon simulations, leads to reduced local membrane curvature that may facilitate the detachment of COPII-coated carriers from the ER membrane. After dissociation from the ER, multiple COPII-coated carriers undergo clustering before homotypic fusion or heterotypic fusion with pre-existing ER-Golgi Intermediate Compartment (ERGIC) membranes, which is facilitated by TFG, a protein that likely undergoes liquid-liquid phase separation (LLPS).(36, 37) To understand how LLPS promotes COPII carrier fusion events, we need to model these proteins at the coarse-grained level, as atomistic simulations will be computationally prohibitive for such problems. Nevertheless, our atomistic simulations of Sar1/membrane interactions serves as a basis to build and calibrate efficient coarse-grained models for various COPII components.

## Methods

We employ AlphaFold2(25) to predict the structures of Sar1 in both GDP-(PDB code 1F6B(8)) and GTP-(PDB code 1M2O(9)) bound states referred to as the h-GDP and y-GTP structure, respectively. We also utilize CHARMM-GUI(38) to generate Sar1 models in both nucleotide-bound states. We build the h-GTP model of Sar1 using the GDP state sequence and the y-GTP structure as a template with the SWISS-MODEL(39) webserver and the Molefracture plugin of VMD(40). Comparison of the various structural models of Sar1 is shown in Fig-S1. (see SI text Sec. 1A for details) We first perform solution simulations of the AlphaFold2 derived models in the absence of membrane. Explicit solvent all-atom MD simulations are performed using GROMACS(41, 42) version 2018.3 and the CHARMM36m(43) forcefield with the TIP3P explicit solvent model. The systems are set up using CHARMM-GUI(38, 44, 45). The protein is placed in a ~ 46 × 46 × 46 Å^3^ box. Na^+^ and Cl^-^ ions are added to neutralize the system and to maintain the physiological (0.15 M) salt concentration. MD trajectories are used for the computation of Cumulative Variance of Coordinated Fluctuations (CVCF)(27) to characterize the flexibility of the amino-terminal region of SAR1 (see SI text Sec. 1B for details). Next, we choose 10 different conformations spanning over 300 ns of the production run and supply them to membrane insertion simulations. To better sample protein insertion and equilibrate lipid distributions around the protein, the protein-membrane simulations are performed in two steps. In the first step, we employ the Highly Mobile Membrane Mimetic (HMMM) model(46) of membrane bilayer with 1.7 times amplified lipid surface area to facilitate accelerated insertion of the protein into the membrane. The composition of the membrane is 66 % DOPC, 21 % DOPE, 8 % DOPS, and 5 % DOPA as described in Hanna et. al.(14). The area of the membrane is kept constant throughout the HMMM simulation using semi-isotropic pressure coupling where compressibility along the *xy* directions is kept at 0 bar^-1^ (see SI text Sec. 1C for details).

In the second step, the organic solvent, 1,1-dichloroethane, in the HMMM membrane is removed and the short lipid tails are regrown to their full length using CHARMM-GUI (see SI text Sec. 1D for details). In this step, we estimate the membrane penetration depth of proteins from their mass density profile where average penetration depth inside membrane is defined as the mean of the peak position and end point of the mass density of the protein amino-terminus. In order to compare the penetration depths of various Sar1 conformations, we normalize the penetration depth with respect to the thickness of the protein bearing leaflet of the membrane.(see SI text Sec. 1E for details) For estimating protein volume inserted (*V_p_*) into the membrane we simply count the number of atoms (*N_p_*) in between the phosphate planes of the membrane. In the case of Sar1 dimers, we normalize *N_p_* with respect to the distance between the geometric center of the constituent monomers *l* (in nm). In case of the h-GTP-bound dimer, *l* is found to be 1.04 nm.

Finally, membrane ribbon(26) simulations (Fig-S8) are performed with the protein membrane system from the previous step as a seed. Multiple replicas of the protein-membrane system along +*X* and −*X* directions are generated using the gmx genbox utility followed by the deletion of membrane segments around the terminus along +*X* and −*X* to discard the possibility of periodic boundary conditions along *X*. Simulation conditions and parameters are similar to those in solution simulation except that anisotropic pressure coupling is used to avoid merging of the membrane ribbon with its periodic images. Off-diagonal compressibility is set to zero to maintain the rectangular shape of the box. Details of the box size, number of atoms, and the timescale of simulations are summarized in SI (Table-S1, see SI text Sec. 1F for details).

## ACKNOWLEDGMENTS

The work is supported in part by the grant NSF-DMS1661900 to AA and QC and NIH R35GM134865 to AA. Computational resources from the Extreme Science and Engineering Discovery Environment (XSEDE(49)), which is supported by NSF grant number ACI-1548562, are greatly appreciated; part of the computational work was performed on the Shared Computing Cluster which is administered by Boston University’s Research Computing Services (URL: www.bu.edu/tech/support/research/).

## Notes

The authors have no competing interests.

### Competing Interest Statement

The authors have declared no competing interest.

## References

1. C Barlowe, et al., Copii: a membrane coat formed by sec proteins that drive vesicle budding from the endoplasmic reticulum. Cell 77, 895–907 (1994).

2. D Jensen, R Schekman, Copii-mediated vesicle formation at a glance. J. Cell Sci. 124, 1–4 (2011).

3. C d’Enfert, LJ Wuestehube, T Lila, R Schekman, Sec12p-dependent membrane binding of the small gtp-binding protein sar1p promotes formation of transport vesicles from the er. J. Cell Biol. 114, 663–670 (1991).

4. MC Lee, et al., Sar1p n-terminal helix initiates membrane curvature and completes the fission of a copii vesicle. Cell 122, 605–617 (2005).

5. X Bi, JD Mancias, J Goldberg, Insights into copii coat nucleation from the structure of sec23* sar1 complexed with the active fragment of sec31. Dev. Cell 13, 635–645 (2007).

6. E Miller, B Antonny, S Hamamoto, R Schekman, Cargo selection into copii vesicles is driven by the sec24p subunit. EMBO J. 21, 6105–6113 (2002).

7. W Ma, J Goldberg, Tango1/ctage5 receptor as a polyvalent template for assembly of large copii coats. Proc. Natl. Acad. Sci. U. S. A. 113, 10061–10066 (2016).

8. M Huang, et al., Crystal structure of sar1-gdp at 1.7 a resolution and the role of the nh2 terminus in er export. J. Cell Biol. 155, 937–948 (2001).

9. X Bi, RA Corpina, J Goldberg, Structure of the sec23/24–sar1 pre-budding complex of the copii vesicle coat. Nature 419, 271–277 (2002).

10. B Antonny, S Beraud-Dufour, P Chardin, M Chabre, N-terminal hydrophobic residues of the g-protein adp-ribosylation factor-1 insert into membrane phospholipids upon gdp to gtp exchange. Biochemistry 36, 4675–4684 (1997).

11. M Franco, P Chardin, M Chabre, S Paris, Myristoylation-facilitated binding of the g protein arf1gdp to membrane phospholipids is required for its activation by a soluble nucleotide exchange factor (). J. Biol. Chem. 271, 1573–1578 (1996).

12. JG Donaldson, CL Jackson, Arf family g proteins and their regulators: roles in membrane transport, development and disease. Nat. Rev. Mol. Cell Biol. 12, 362–375 (2011).

13. P Chavrier, J Ménétrey, Toward a structural understanding of arf family: effector specificity. Structure 18, 1552–1558 (2010).

14. MG Hanna, et al., Sar1 gtpase activity is regulated by membrane curvature. J. Biol. Chem. 291, 1014–1027 (2016).

15. GS Ayton, GA Voth, Multiscale simulation of protein mediated membrane remodeling in Seminars in cell & developmental biology. (Elsevier), Vol. 21, pp. 357–362 (2010).

16. H Yu, K Schulten, Membrane sculpting by f-bar domains studied by molecular dynamics simulations. PLoS Comput. Biol. 9, e1002892 (2013).

17. A Arkhipov, Y Yin, K Schulten, Four-scale description of membrane sculpting by bar domains. Biophys. J. 95, 2806–2821 (2008).

18. A Arkhipov, Y Yin, K Schulten, Membrane-bending mechanism of amphiphysin n-bar domains. Biophys. J. 97, 2727–2735 (2009).

19. PD Blood, GA Voth, Direct observation of bin/amphiphysin/rvs (bar) domain-induced membrane curvature by means of molecular dynamics simulations. Proc. Natl. Acad. Sci. U. S. A. 103, 15068–15072 (2006).

20. PD Blood, RD Swenson, GA Voth, Factors influencing local membrane curvature induction by n-bar domains as revealed by molecular dynamics simulations. Biophys. J. 95, 1866–1876 (2008).

21. T Mandal, W Lough, SE Spagnolie, A Audhya, Q Cui, Molecular simulation of mechanical properties and membrane activities of the escrt-iii complexes. Biophys. J. 118, 1333–1343 (2020).

22. T Mandal, SE Spagnolie, A Audhya, Q Cui, Protein-induced membrane curvature in coarsegrained simulations. Biophys. J. 120, 3211–3221 (2021).

23. L Harker-Kirschneck, et al., Physical mechanisms of escrt-iii–driven cell division. Proc. Natl. Acad. Sci. U. S. A. 119 (2022).

24. AJ Sodt, RW Pastor, Molecular modeling of lipid membrane curvature induction by a peptide: more than simply shape. Biophys. J. 106, 1958–1969 (2014).

25. J Jumper, et al., Highly accurate protein structure prediction with alphafold. Nature 596, 583–589 (2021).

26. Z Wu, K Schulten, Synaptotagmin’s role in neurotransmitter release likely involves ca2+-induced conformational transition. Biophys. J. 107, 1156–1166 (2014).

27. S Paul, SRK Ainavarapu, R Venkatramani, Variance of atomic coordinates as a dynamical metric to distinguish proteins and protein–protein interactions in molecular dynamics simulations. J. Phys. Chem. B 124, 4247–4262 (2020).

28. H Hariri, N Bhattacharya, K Johnson, AJ Noble, SM Stagg, Insights into the mechanisms of membrane curvature and vesicle scission by the small gtpase sar1 in the early secretory pathway. J. Mol. Biol. 426, 3811–3826 (2014).

29. KR Long, et al., Sar1 assembly regulates membrane constriction and er export. J Cell Biol 190, 115–128 (2010).

30. A Bielli, et al., Regulation of sar1 nh2 terminus by gtp binding and hydrolysis promotes membrane deformation to control copii vesicle fission. J. Cell Biol. 171, 919–924 (2005).

31. J Hutchings, V Stancheva, EA Miller, G Zanetti, Subtomogram averaging of copii assemblies reveals how coat organization dictates membrane shape. Nat. Commun. 9, 1–8 (2018).

32. K Matsuoka, et al., Copii-coated vesicle formation reconstituted with purified coat proteins and chemically defined liposomes. Cell 93, 263–275 (1998).

33. F Adolf, et al., Scission of copi and copii vesicles is independent of gtp hydrolysis. Traffic 14, 922–932 (2013).

34. AF Loftus, VL Hsieh, R Parthasarathy, Modulation of membrane rigidity by the human vesicle trafficking proteins sar1a and sar1b. Biochem. Biophys. Res. Commun. 426, 585–589 (2012).

35. F Campelo, HT McMahon, MM Kozlov, The hydrophobic insertion mechanism of membrane curvature generation by proteins. Biophys. J. 95, 2325–2339 (2008).

36. YG Zhao, H Zhang, Phase separation in membrane biology: the interplay between membranebound organelles and membraneless condensates. Dev. Cell 55, 30–44 (2020).

37. MG Hanna, et al., Tfg facilitates outer coat disassembly on copii transport carriers to promote tethering and fusion with er–golgi intermediate compartments. Proc. Natl. Acad. Sci. U. S. A. 114, E7707–E7716 (2017).

38. S Jo, T Kim, VG Iyer, W Im, Charmm-gui: a web-based graphical user interface for charmm. J. Comput. Chem. 29, 1859–1865 (2008).

39. A Waterhouse, et al., Swiss-model: homology modelling of protein structures and complexes. Nucleic Acids Res. 46, W296–W303 (2018).

40. W Humphrey, A Dalke, K Schulten, VMD – Visual Molecular Dynamics. J Mol Graph 14, 33–38 (1996).

41. MJ Abraham, et al., Gromacs: High performance molecular simulations through multi-level parallelism from laptops to supercomputers. SoftwareX 1, 19–25 (2015).

42. HJ Berendsen, D van der Spoel, R van Drunen, Gromacs: A message-passing parallel molecular dynamics implementation. Comput. Phys. Commun. 91, 43–56 (1995).

43. J Huang, et al., Charmm36m: an improved force field for folded and intrinsically disordered proteins. Nat. Methods 14, 71–73 (2017).

44. BR Brooks, et al., Charmm: the biomolecular simulation program. J. Comput. Chem. 30, 1545–1614 (2009).

45. J Lee, et al., Charmm-gui input generator for namd, gromacs, amber, openmm, and charmm/openmm simulations using the charmm36 additive force field. J. Chem. Theory Comput. 12, 405–413 (2016).

46. YZ Ohkubo, TV Pogorelov, MJ Arcario, GA Christensen, E Tajkhorshid, Accelerating membrane insertion of peripheral proteins with a novel membrane mimetic model. Biophys. J. 102, 2130–2139 (2012).

