## Supplementary Information for "Molecular mechanism of GTP binding- and dimerization-induced enhancement of Sar1-mediated membrane remodeling"

#### This PDF file includes:

Supplementary text
Figs. S1 to S12
Table S1
SI References

### Supporting Information Text

### 1. Methods

**A. Modeling.** The crystal structures of Sar1 are available in both GDP and GNP (GTP) bound states, with the PDB code 1F6B(1) and 1M2O(2) respectively (Fig-S1A-B); they were for Sar1 from hamster (*Cricetulus griseus*) and yeast (*Saccharomyces cerevisiae*), respectively. Apart from the missing amino-terminal region as mentioned above, coordinates of few other residues (residues 49-54 and 79-82 in the GDP structure and residues 157-159 in the GTP structure) were also unresolved. We utilize the v2.0.0 version of alphafold (AlphaFold2)(3) to predict the structures of Sar1 in both GTP and GDP bound states using the sequences of the corresponding crystal structures, and the resulting models are referred to as the h-GDP and y-GTP structure, respectively, in the discussion; the missing amino-terminal helix was predicted to be helical in both nucleotide states (Fig-S1C-D). Apart from the AlphaFold2 based structures, we also employ CHARMM-GUI(4) to model the missing coordinates; the amino-terminus mostly acquires a random coil structure in those models (Fig-S1E-F). Finally, we also build a GTP state model for the hamster sequence using the y-GTP structure as a template with the SWISS-MODEL(5) webserver, which succeeded in building the structural model for residues 7-196. We then generate a helical structure for residues 1-6 and a coil segment for residues 196-198 using the Molefracture plugin of VMD(6) and add these segments to the SWISS-MODEL derived structure; finally, GTP is added to the binding pocket, completing the h-GTP model. Comparison of the various structural models of Sar1 yields RMSD values in the range of 1.9-6.6 Å (Fig-S1).

**B. Protein simulation in the absence of membrane.** We first perform solution simulations of the AlphaFold2 derived models in the absence of membrane. All explicit solvent all-atom MD simulations are performed using GROMACS(7, 8) version 2018.3 and the CHARMM36m(9) forcefield with the TIP3P explicit solvent model. The systems are set up using CHARMM-GUI(4, 10, 11). The protein is placed in a $\sim 46 \times 46 \times 46$ Å$^3$ box. $Na^+$ and $Cl^-$ ions are added to neutralize the system and to maintain the physiological (0.15 M) salt concentration. Periodic boundary conditions are employed along all three principal directions. The Particle Mesh Ewald(12) (PME) method is used to compute the electrostatic interactions and a switching function is used to reduce the van der Waals force smoothly to zero between 1.0 and 1.2 nm. The solvated system is first energy minimized using the conjugate gradient method to remove bad contacts between the solute and solvent atoms. This step is followed by a short NVT simulation in which a harmonic restraint is initially applied to the protein atoms and then gradually released during equilibration. This NVT-equilibrated system is then subjected to NPT production run for 300 ns at the atmospheric pressure and 303 K temperature, during which no atoms are restrained. The temperature of the system during equilibration is controlled by the Nosé-Hoover thermostat(13, 14) with a time constant of 1 ps. Additionally, Parrinello-Rahman barostat(15) with a time constant of 5 ps is employed during the production run to maintain the pressure of the system to 1 bar. The LINCS algorithm(16) is used to constrain covalent bonds involving hydrogen atoms to enable an integration time step of 2 fs. MD trajectories are used for the computation of Cumulative Variance of Coordinated Fluctuations (CVCF)(17) to characterize the flexibility of the amino-terminal region of Sar1. CVCF is defined as follows

$$\sigma^2_{CVCF} = \sum_{i=1}^{N} \frac{1}{T_S} \sum_{t=1}^{T_S} [x_i(t) - \frac{1}{T_S} \sum_{t'=1}^{T_S} x_i(t')]^2 \qquad [1]$$

where $N$ represents the set of backbone atoms of the amino-terminal segment (resid 1-12 and 13-23), $T_S$ goes up to $\sim$ 300 ns and $x_i$ is the coordinate of $i$th atom at time $t$. During the CVCF analysis, rotation and translation of the protein molecule is eliminated by aligning all backbone atoms of the protein from each frame with respect to that from the first frame.

**C. HMMM membrane simulation.** Next, we choose 10 different conformations spanning over 300 ns of the production run and supply them to membrane insertion simulations. To better sample protein insertion and equilibrate lipid distributions around the protein, the protein-membrane simulations are performed in two steps. In the first step, we employ the Highly Mobile Membrane Mimetic (HMMM) model(18) of membrane bilayer with 1.7 times amplified lipid surface area to facilitate accelerated insertion of the protein into the membrane. The composition of the membrane is 66 % DOPC, 21 % DOPE, 8 % DOPS, and 5 % DOPA as described in Hanna et. al.(19). We do not include any cholesterol in our membrane as our HMMM model already leads to sufficient protein binding. Using the CHARMM-GUI HMMM builder(20, 21), we build the initial model of protein-membrane system where the protein is placed at varying distances from the membrane while keeping the amino-terminal helix oriented on top of the membrane. Energy minimization is followed by a short NPT equilibration that gradually releases harmonic constraints on heavy atoms; finally NPT production run of timescale $\sim$ 80 to 160 ns is performed for each protein-membrane system. The pressure and temperature of the system during this step are controlled by the Berendsen barostat(22) with a time constant of 5 ps and the Berendsen thermostat(22) with a time constant of 1 ps, respectively. The area of the membrane is kept constant throughout the HMMM simulation using semiisotropic pressure coupling where compressibility along the $xy$ directions is kept at 0 bar$^{-1}$.

**D. Full lipid model.** In the second step, the organic solvent, 1,1-dichloroethane, in the HMMM membrane is removed and the short lipid tails are regrown to their full length using CHARMM-GUI. Subsequently, NPT simulations are carried out for 160 ns with the Nosé-Hoover thermostat and Parinello-Rahman barostat to control temperature and pressure of the system, respectively; compressibility is turned on along the $xy$ direction in semiisotropic pressure coupling. Trajectories obtained in this step are used for the assessment of mass density profile and protein membrane contact using the gmx density and gmx mindist (cutoff: 5 Å) utilities of GROMACS, respectively.

**E. Penetration depth analysis.** We estimate the membrane penetration depth of proteins from their mass density profile where phosphate plane and the C318 atoms are considered as the periphery and the core of the membrane respectively. Define $d_1$ as the inter-peak distance of the mass density distribution of the phosphate group of the DOPC atoms of the membrane and the N terminus of the protein. Next, $d_2$ is defined as the distance where the amino-terminal mass density decays to 0 with respect to the phosphate plane of the membrane. We estimate the average penetration depth of proteins inside membrane $d$ as the mean of $d_1$ and $d_2$. The thickness of the protein bearing leaflet of the membrane $d_m$ is defined as the inter-peak distance of the mass density distributions of periphery (phosphate) and core (C318) of the membrane. In order to compare the penetration depths of various Sar1 conformations, we normalize the penetration depth with respect to the membrane leaflet thickness $((d/d_m) * 100)$.

**F. Analysis of protein volume inclusion into the membrane.** We estimate the amount of protein volume ($V_p$) inside membrane by simply counting the number of atoms ($N_p$) between the phosphate planes of the membrane. In each frame of 50-160 ns simulation, we calculate the number of atoms ($N_p$) that lie between the geometric centers of the upper and lower phosphate planes. An average and standard deviation of $N_p$ is computed over the frames. A greater $N_p$ value indicates a larger $V_p$. Naturally, this procedure may yield a higher $V_p$ in case of a dimer compared to the case of a monomer. Thus we estimate $V_p$ by normalizing $N_p$ with respect to the separation ($l$, in nm) between the constituent protein fragments in a protein multimer; i.e., $V_p = N_p/l$, where $l = 1.04$ nm for the case of the h-GTP dimer.

**G. Membrane ribbon simulation.** Finally, membrane ribbon(23) simulations (Fig-S8) are performed with the protein membrane system from the previous step as a seed. Multiple replicas of the protein-membrane system along $+X$ and $-X$ directions are generated using the gmx genbox utility followed by the deletion of membrane segments around the terminus along $+X$ and $-X$ to discard the possibility of periodic boundary conditions along $X$. Simulation conditions and parameters are similar to those in solution simulation except that anisotropic pressure coupling is used to avoid merging of the membrane ribbon with its periodic images. Off-diagonal compressibility is set to zero to maintain the rectangular shape of the box. Details of the box size, number of atoms, and the timescale of simulations are summarized in Table-S1.

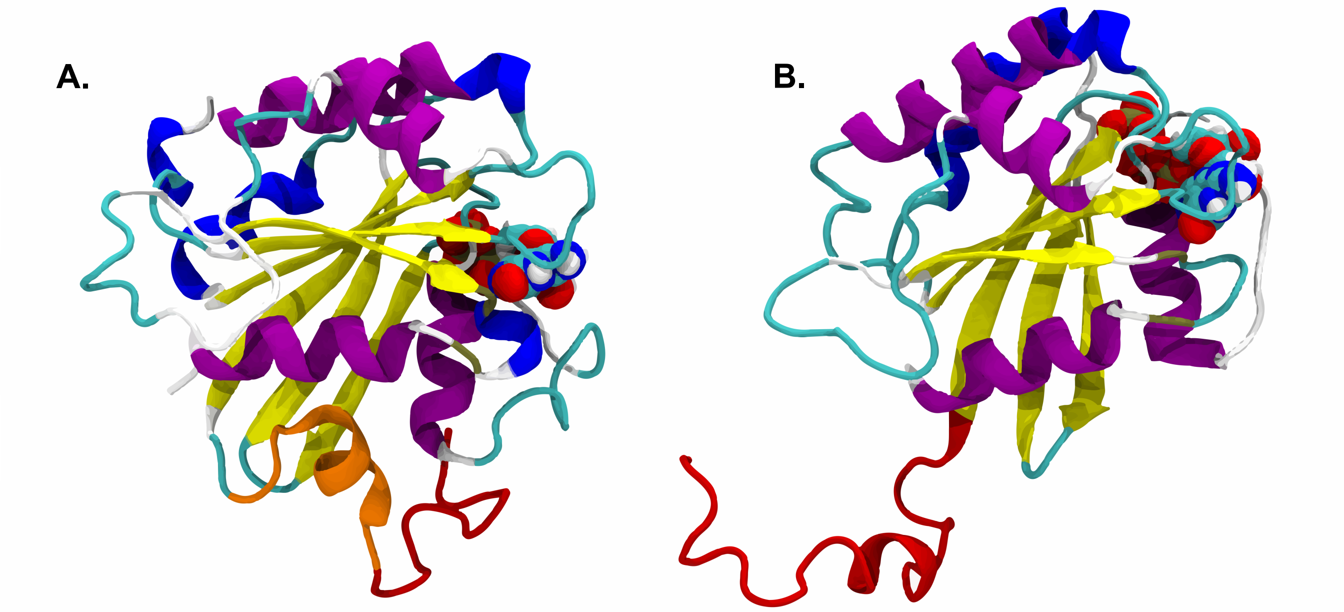


**Fig. S1.** CHARMM-GUI derived model of (A) GDP- and (B) GTP-bound Sar1. GDP and GTP are shown in the van der Waals representation. The amino-terminal residues 1-12 in the GDP-bound and residues 1-23 in the GTP-bound Sar1 are modelled (highlighted in red color) using CHARMM-GUI since their coordinates were not resolved in the corresponding crystal structures. These modelled regions remain unstructured unlike the AlphaFold2 derived models. Coordinates of residues 13-23 were resolved in the crystal structure of GDP-bound Sar1 and the region is highlighted in orange.

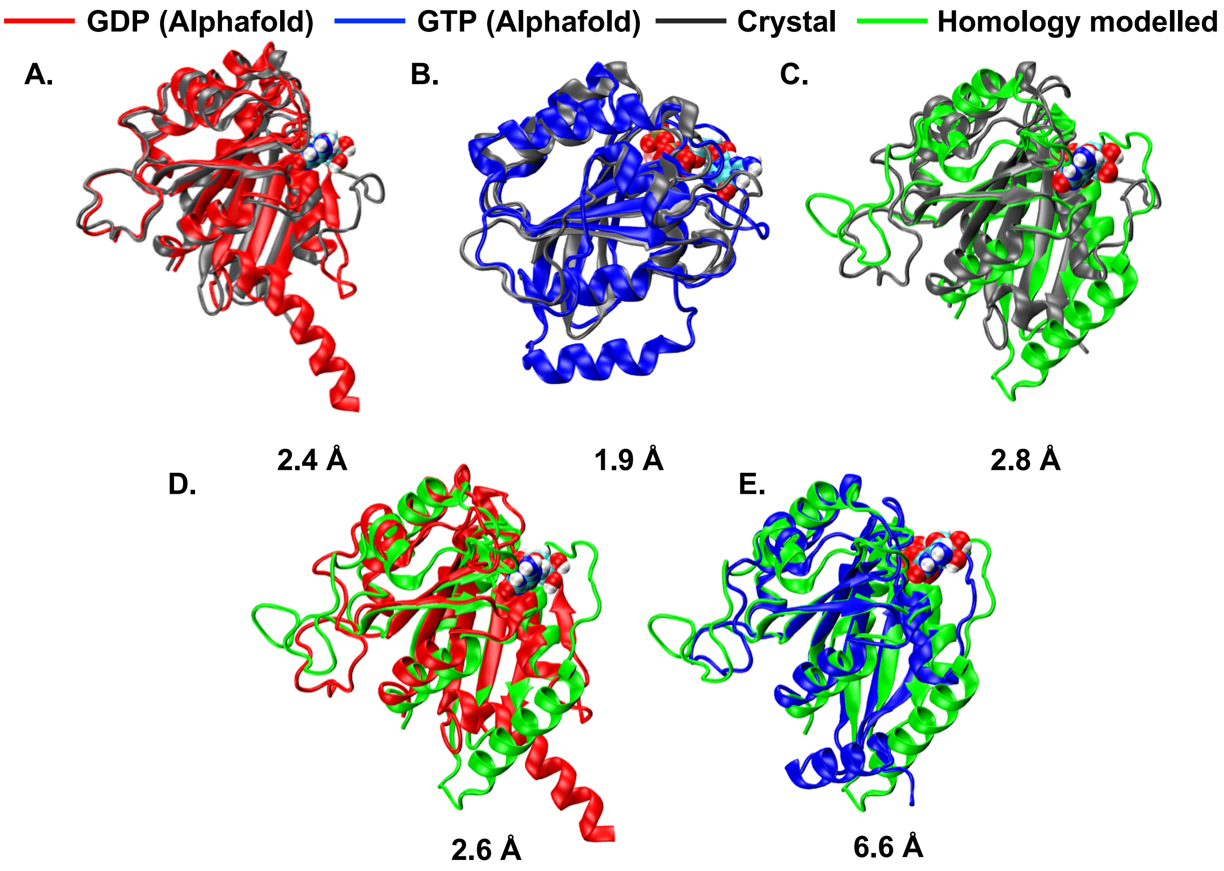


**Fig. S2.** Alignment of different models of Sar1. (A-B) Alignment of Alphafold2 model of GDP- and GTP-bound states with respect to the corresponding crystal structures yield 2.4 Å and 1.9 Å RMSD respectively. Homology modelled Sar1 aligned with the (C) GDP-bound crystal structure, (D) Alphafold modelled GDP-bound state and (E) Alphafold modelled GTP-bound state exhibit RMSD of 2.8 Å, 2.6 Å and 6.6 Å respectively. The RMSD values are computed for backbone atoms excluding the amino terminus (residues 1-23) and the loop region (48-62 and 152-172).

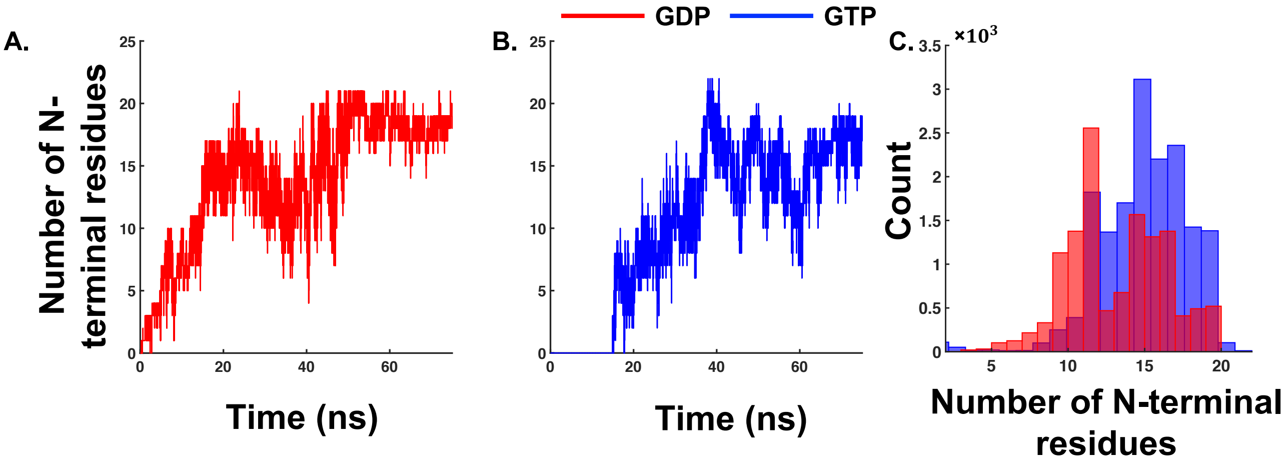


**Fig. S3.** Assessment of insertion of the AlphaFold2 derived Sar1 model into the HMMM membrane. The number of amino-terminal residues inserted into the membrane as simulation progresses in a representative trajectory of (A) GDP- and (B) GTP-bound Sar1. If C$\alpha$ atom of any amino-terminal residue (residue 1-23) in a frame lies between the centers of mass of phosphate atoms of the upper and lower leaflets of the membrane, then the residue is counted as inserted into the membrane in that frame. (C) Histogram of the number of amino-terminal residues inserted into the membrane from 10 different MD trajectories of GDP- (red) and GTP- (blue) bound Sar1. 10 trajectories originate from different initial conformations of the protein obtained from solution simulations.

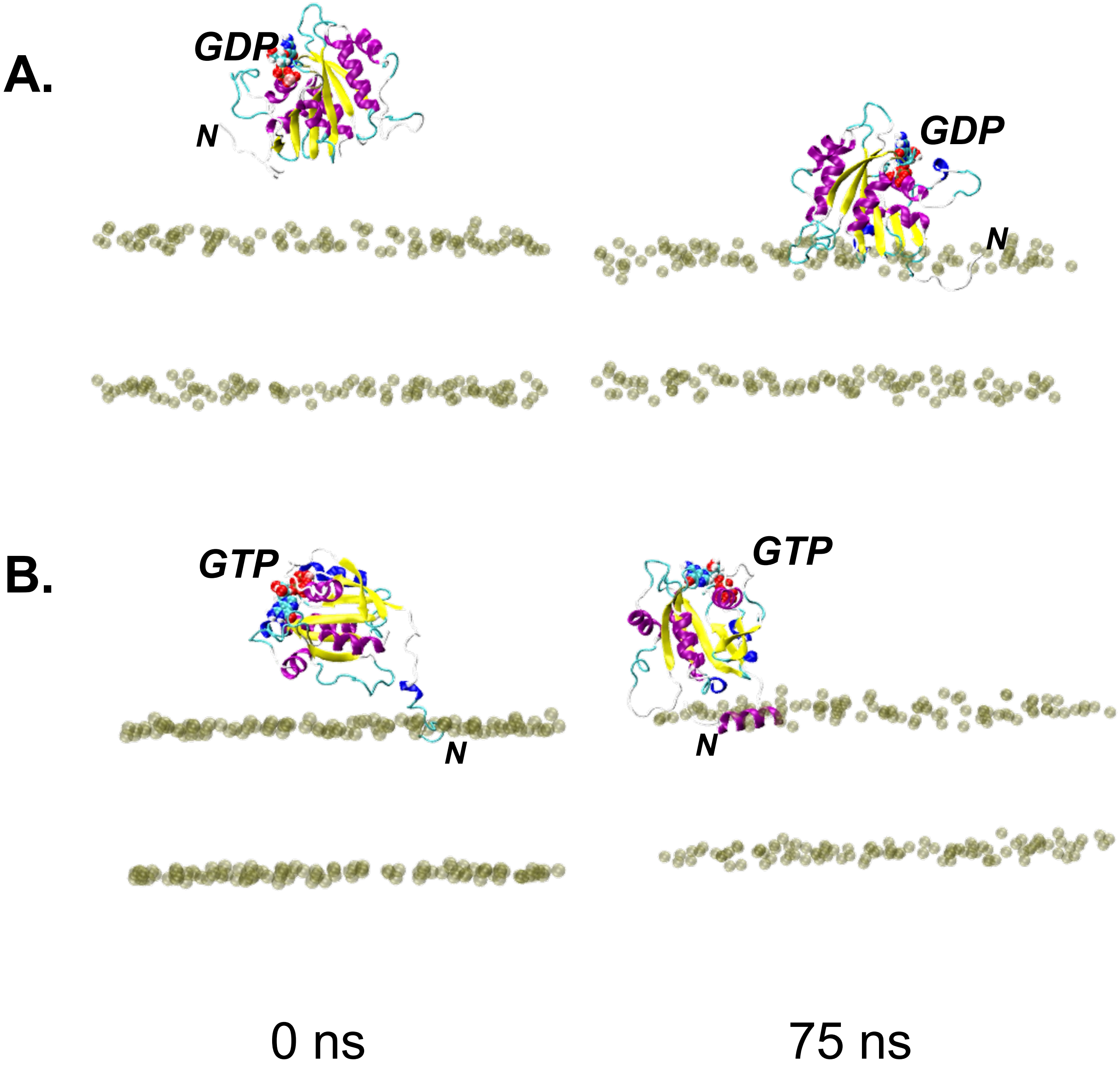


**Fig. S4.** Membrane induced helicity in the amino-terminus of the GTP-bound Sar1. Snapshots of Sar1 in which the unresolved amino-terminus is modelled as a random coil using CHARMM-GUI at 0 ns (left) and 75 ns (right) for the (A) GDP- and (B) GTP-bound state in the presence of the HMMM membrane. The amino-terminal region remains in the coil state after inserting into the membrane in the GDP-bound state. By contrast, the amino-terminus of the GTP-bound state adopts the $\alpha$-helical structure after penetrating into the membrane. Only P atoms of the phosphate groups in the lipids are shown for clarity.

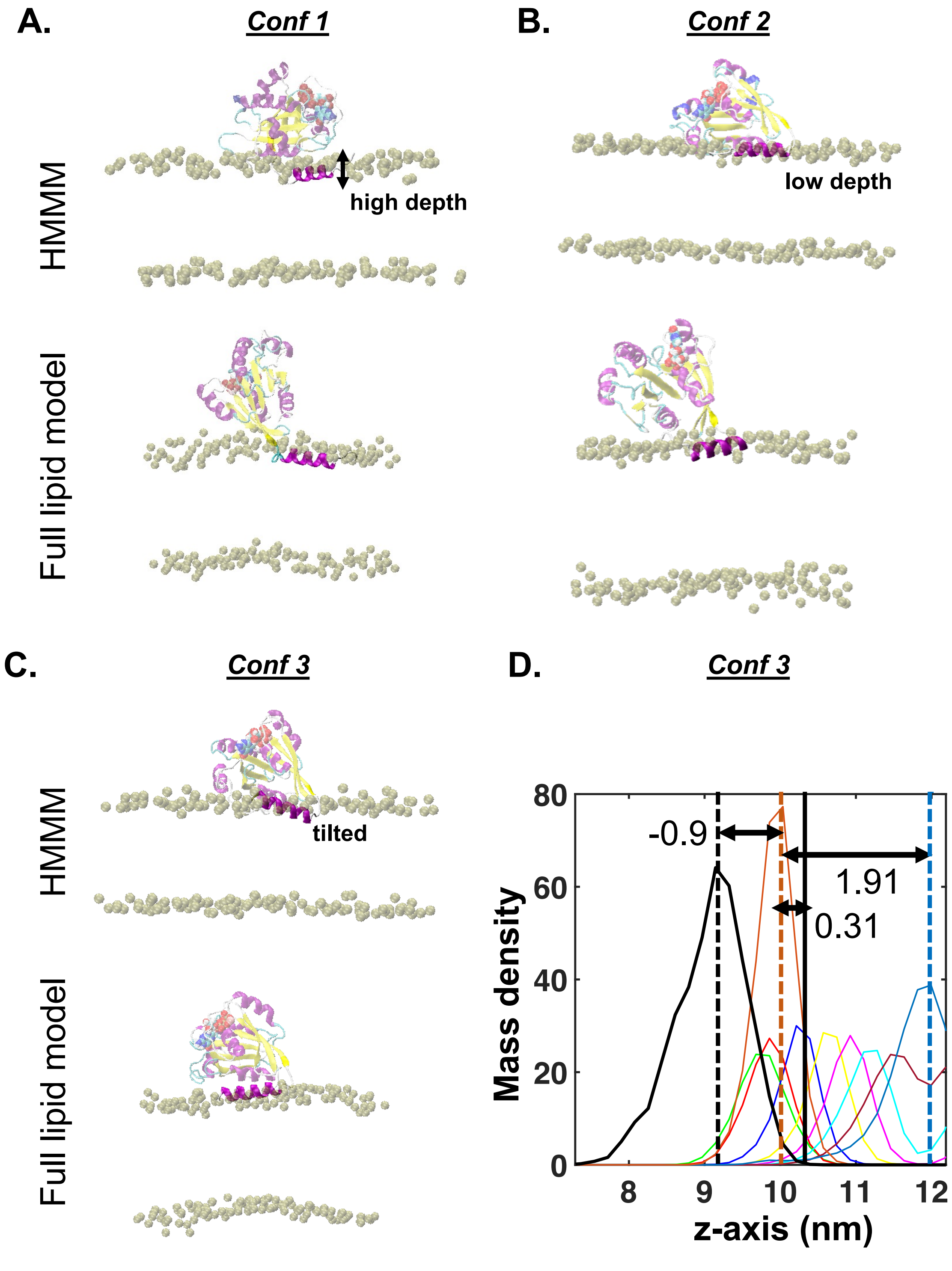


**Fig. S5.** Examining the ability of various amino-terminal conformations of the GTP bound Sar1 to insert into the membrane with full lipid tail. (A-C) Snapshots of protein membrane interface at the end of the simulation with the HMMM membrane model (top) and full lipid model (down) for 3 different conformations. (D) Mass density profile of various atoms of the DOPC lipid and the protein amino-terminus (black) are shown for conformation 3. DOPC lipid atoms are color-coded as shown in Fig-3 of the main text. Conformation 1 exhibits a higher penetration compared to conformation 2 in the HMMM membrane. As a result, conformation 1 yields a higher penetration depth in the full lipid model as well. The amino-terminus of conformation 3 adopts a tilted conformation, which causes a poor insertion into the full lipid membrane.

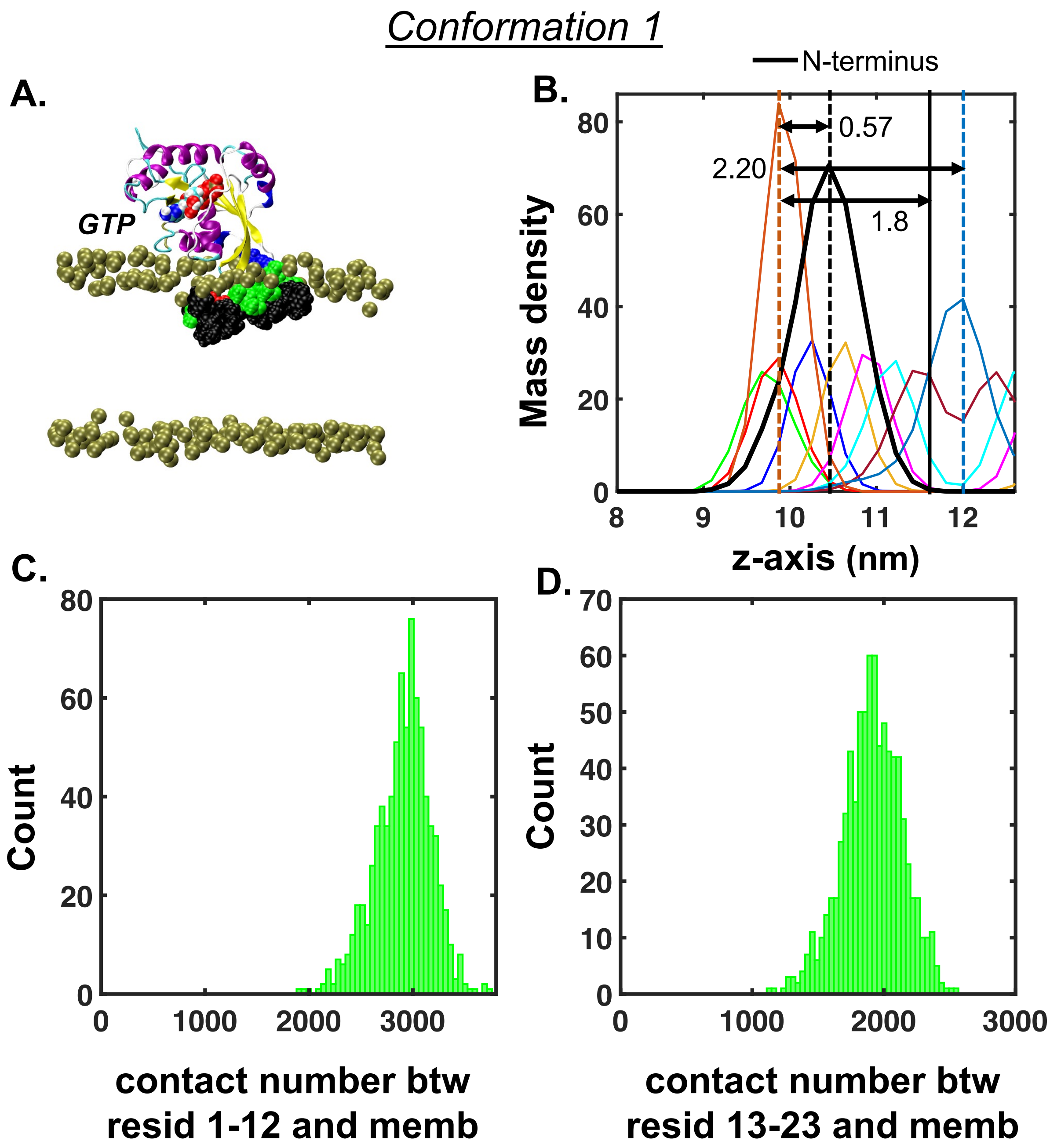


**Fig. S6.** Membrane binding activity of conformation 1 of the homology model of the GTP-bound hamster Sar1 (h-GTP). (A) Protein/membrane interface of h-GTP following 160 ns of simulation. (B) Mass density plots of the amino-terminus of h-GTP Sar1 along with various DOPC atoms indicate that h-GTP penetrates into the membrane to a similar extent as y-GTP (Fig-4C in the main text). Histograms of the numbers of contacts between the membrane and (C) residues 1-12 or (D) residues 13-23 of Sar1.

### Conformation 2

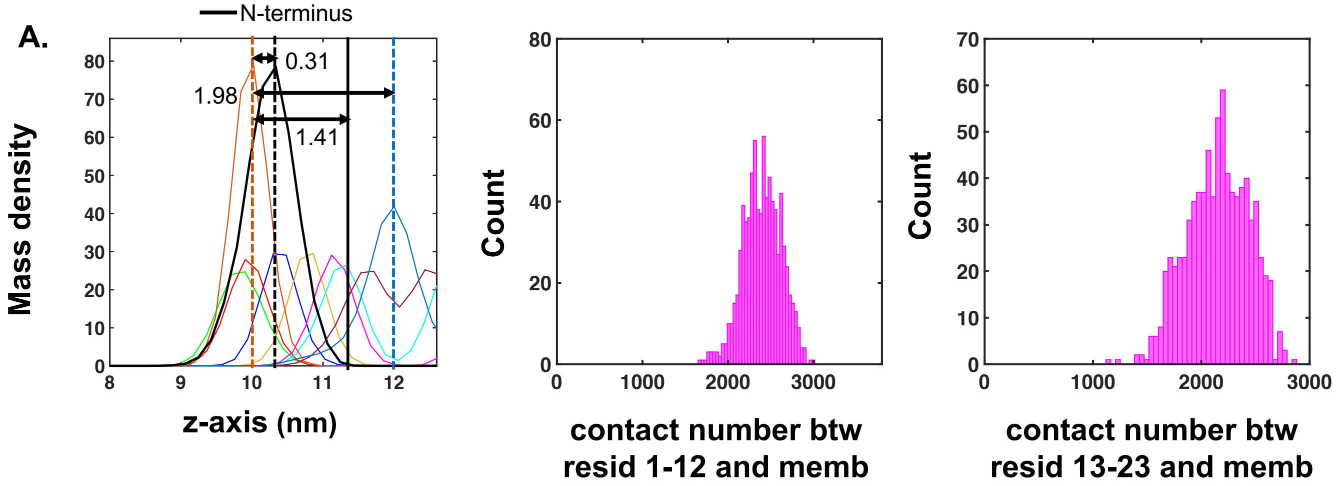


**Fig. S7.** Membrane binding activity of conformation 2 of the homology model of the GTP-bound hamster Sar1 (h-GTP).(A) Mass density plots of the amino-terminus of h-GTP Sar1 along with various DOPC atoms indicate that h-GTP penetrates into the membrane to a similar extent as y-GTP (Fig-4C of the main text). Histograms of the numbers of contacts between the membrane and (C) residues 1-12 or (D) residues 13-23 of Sar1.

 

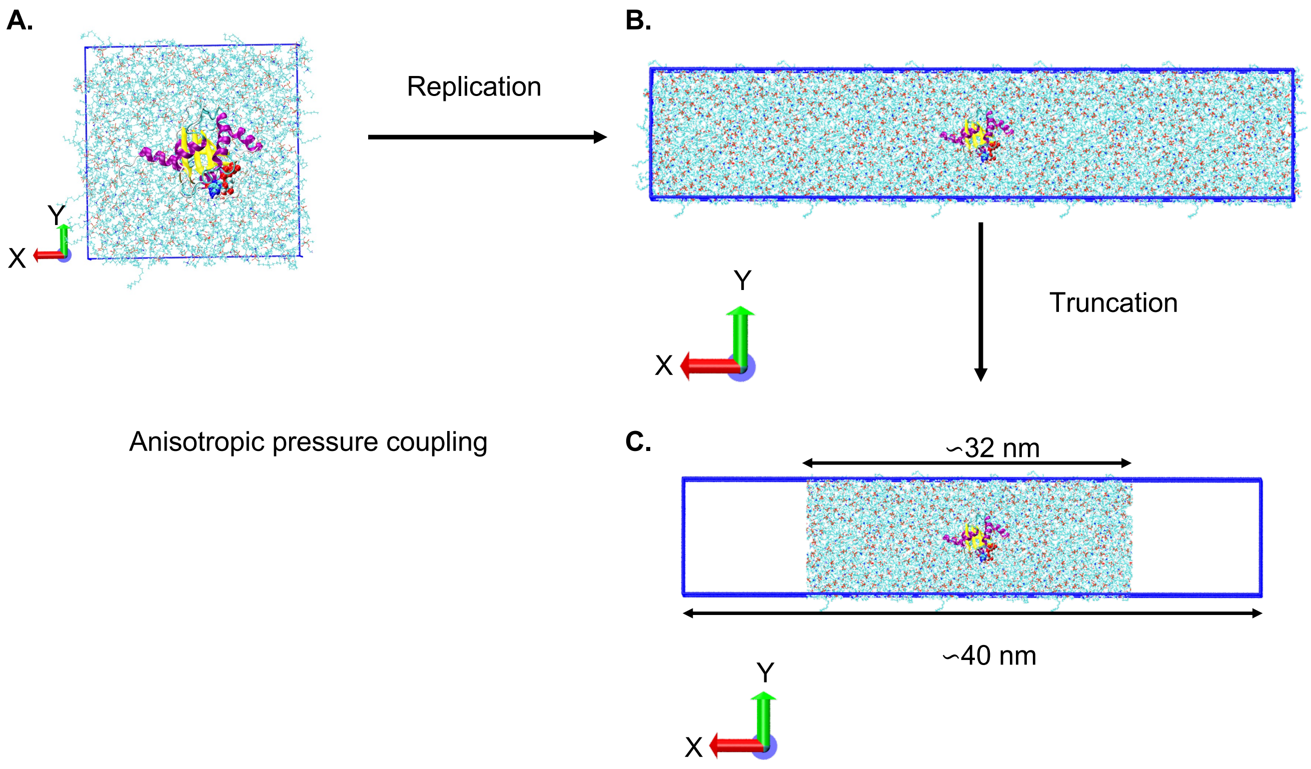


**Fig. S8.** Membrane ribbon simulation protocol. (A) Snapshot of GTP-bound Sar1 inserted into the membrane with the full lipid tail in a $\sim$ 10.9$\times$10.9$\times$11.5 $nm^3$ box (No. 5 in table S1). Snapshot of the same system (B) when the membrane is replicated twice along the $X$ axis and (C) when some lipid molecules are removed from both ends to break the periodicity along the $X$-axis. The simulation box becomes 40 nm in length where the membrane covers 32 nm along the $X$ dimension. The membrane remain periodic only along the $Y$ axis. This protein membrane system is then supplied for simulations (No. 7 in table S1) where the anisotropic pressure coupling is used for non-zero diagonal but zero off-diagonal compressibility.

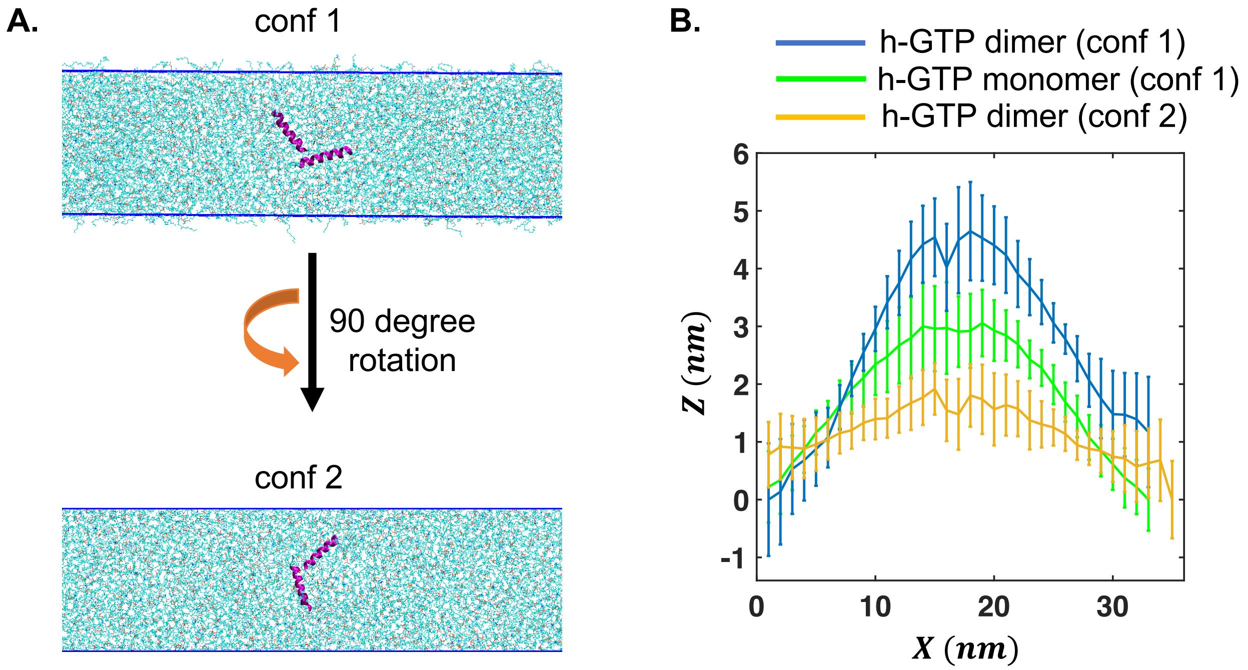


**Fig. S9.** Orientation of the amphipathic helix on the membrane ribbon influences the amount of curvature induced on it. (A) Conformations 1 (Fig 7-8 in the main text) and 2 of the h-GTP dimer where the later is obtained from the former upon 90$^{\circ}$ rotation around the membrane normal. Note that simulations are carried out with the full Sar1 dimer in both orientations but only the amino-terminal helices are shown for clarity. These snapshots illustrate that conformation 1 has a higher volume inclusion projected along the longer (non-periodic) membrane axis than that of conformation 2. (B) The $Z$-positions of the membrane mid-plane averaged over the $Y$-direction as a function of the $X$ position for the membrane ribbon simulations with conformation 1 and 2 of the h-GTP dimer and conformation 1 of the h-GTP monomer. Conformation 2 of the h-GTP dimer induces a much lower positive curvature compared to the conformation 1 of the h-GTP dimer, primarily due to the significantly smaller amount of volume inclusion projected along the non-periodic axis of the membrane ribbon.

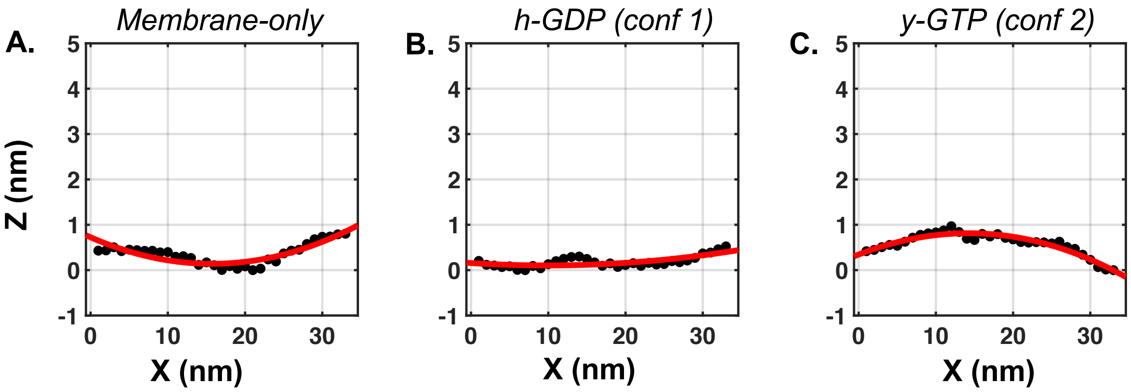


E.

| | Membrane-only | h-GDP (conf 1) | y-GTP (conf 2) |
|---|---|---|---|
| Equation | $y = a - \sqrt{r^2 - (x-b)^2}$ | $y = a - \sqrt{r^2 - (x-b)^2}$ | $y = a + \sqrt{r^2 - (x-b)^2}$ |
| a ($nm$) | 211.4 (166.2, 256.6) | 912 (270.9, 1553) | -214.8 (-244.7, -185) |
| b ($nm$) | 15.7 (14.75, 16.66) | 9.805 (3.928, 15.68) | 14.22 (13.52, 14.92) |
| r ($nm$) | 211.3 (166.1, 256.4) | 911.9 (270.8, 1553) | 215.6 (185.8, 245.4) |
| curvature ($nm^{-1}$) | 0.005 | 0.0011 | 0.0046 |
| Rsq | 0.7683 | 0.5206 | 0.9118 |

**Fig. S10.** Estimation of radius of curvature from membrane ribbon simulations. Black dots are the mean $Z$ coordinates (Fig-7C of the main text) of P atoms of the membrane loaded with (A) no protein (B) conformation 1 of h-GDP, and (C) conformation 2 of y-GTP. Fitted circle (top equation) is shown as the solid red line. (D) Parameters obtained from the fit is tabulated. $r$ is the radius of the circle which determines the curvature. Rsq values indicate the goodness of the fit. Upper and lower limit (95% confidence bound) of each of the fitted parameters is shown inside the parentheses.

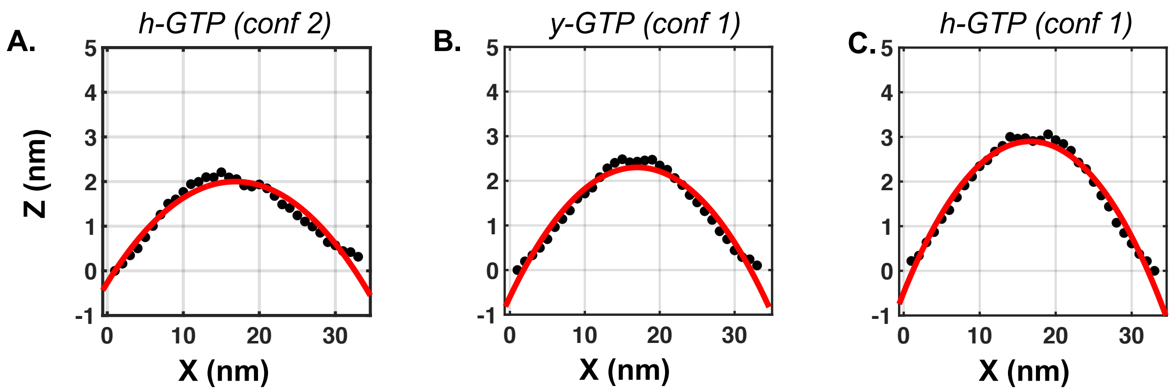


E.

| | h-GTP (conf 2) | y-GTP (conf 1) | h-GTP (conf 1) |
|---|---|---|---|
| Equation | $y = a + \sqrt{r^2 - (x - b)^2}$ | $y = a + \sqrt{r^2 - (x - b)^2}$ | $y = a + \sqrt{r^2 - (x - b)^2}$ |
| a ($nm$) | -61.39 (-67.21, -55.56) | -48.83 (-52.58, -45.09) | -39.81 (-42.16, -37.46) |
| b ($nm$) | 16.8 (16.4, 17.19) | 16.98 (16.66, 17.3) | 16.66 (16.42, 16.9) |
| r ($nm$) | 63.38 (57.63, 69.14) | 51.13 (47.45, 54.81) | 42.71 (40.42, 45) |
| curvature ($nm^{-1}$) | 0.0158 | 0.019 | 0.023 |
| Rsq | 0.941 | 0.9609 | 0.9771 |

**Fig. S11.** Estimation of radius of curvature from membrane ribbon simulations. Black dots are the mean $Z$ coordinates (Fig-7C of the main text) of P atoms of the membrane loaded with (A) conformation 2 of h-GTP (B) conformation 1 of y-GTP, and (C) conformation 1 of h-GTP. Fitted circle (top equation) is shown as the solid red line. (D) Parameters obtained from the fit is tabulated. $r$ is the radius of the circle which determines the curvature. Rsq values indicate the goodness of the fit. Upper and lower limit (95% confidence bound) of each of the fitted parameters is shown inside the parentheses.

A.

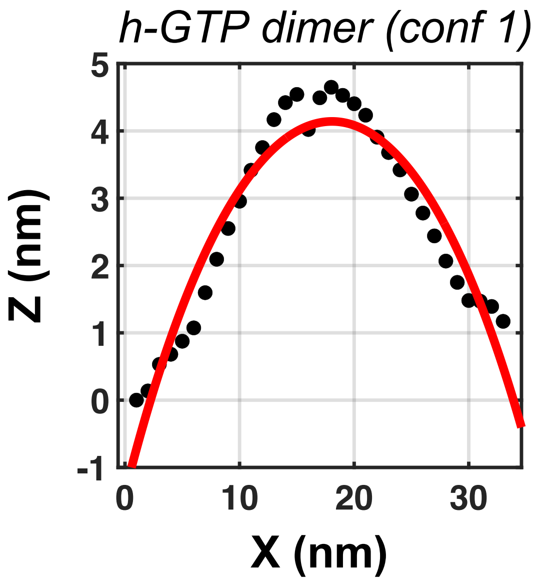


B.

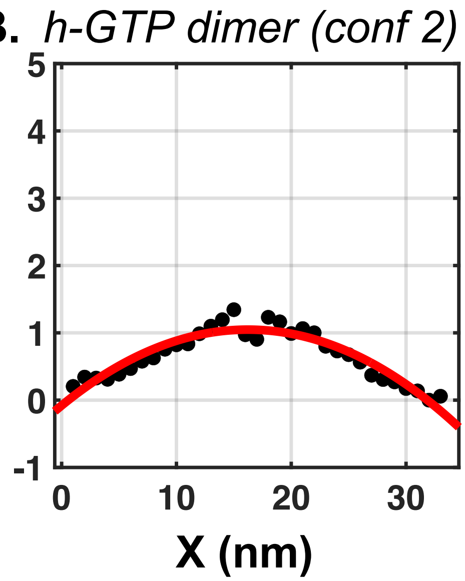


C.

| | h-GTP dimer (conf 1) | h-GTP dimer (conf 2) |
|---|---|---|
| Equation | $y = a + \sqrt{r^2 - (x - b)^2}$ | $y = a + \sqrt{r^2 - (x - b)^2}$ |
| a ($nm$) | -28.2 (-31.59, -24.82) | -116.2 (-131.7, -100.8) |
| b ($nm$) | 18.07 (17.58, 18.55) | 16.22 (15.65, 16.79) |
| r ($nm$) | 32.35 (29.12, 35.57) | 117.3 (101.9, 132.7) |
| curvature ($nm^{-1}$) | 0.0309 | 0.008 |
| Rsq | 0.922 | 0.891 |

**Fig. S12.** Estimation of radius of curvature from membrane ribbon simulations. Black dots are the mean $Z$ coordinates (Fig-7C of the main text) of P atoms of the membrane loaded with (A) conformation 1 and (B) conformation 2 of h-GTP dimer, and (C) conformation 1 of h-GTP. Fitted circle (top equation) is shown as the solid red line. (C) Parameters obtained from the fit is tabulated. $r$ is the radius of the circle which determines the curvature. Rsq values indicate the goodness of the fit. Upper and lower limit (95% confidence bound) of each of the fitted parameters is shown inside the parentheses.

**Table S1. Summary of MD simulations conducted for Sar1 models and their interaction with lipid membrane**

| No. | System<br>(X×Y×Z $nm^3$) | Number of lipids | Total number of atoms | Simulation time (ns) |
|---|---|---|---|---|
| 1 | Sar1 in solution<br>(8.8×8.8×8.8)<br>6 models | - | 70531 | 270 |
| 2 | Sar1 membrane HMMM model<br>(10.9×10.9×12.1)<br>20 conformations | 200 | 120084 | 80 |
| 3 | CHARMM-GUI derived Sar1<br>membrane HMMM model<br>(10.8×10.8×13.5)<br>1 conformation | 200 | 132265 | 75 |
| 4 | Homology modelled Sar1 membrane<br>HMMM model<br>(10.5×10.5×11.8)<br>1 conformation | 200 | 121828 | 75 |
| 5 | Sar1 membrane full lipid model<br>(10.9×10.9×11.5)<br>(5 conformations) | 200 | 118006 | 160 |
| 6 | Membrane curvature simulation<br>without protein<br>(40.6×8×17.1) | 774 | 569963 | 100 |
| 7 | Membrane curvature simulation<br>with protein monomer<br>(40.6×7.9×17.2)<br>5 conformations | 774 | 570073 | 100 |
| 7 | Membrane curvature simulation<br>with protein dimer<br>(40.1×8.1×16.1)<br>2 conformations | 774 | 552183 | 100 |

### References


1. M Huang, et al., Crystal structure of sar1-gdp at 1.7 a resolution and the role of the nh2 terminus in er export. *J. Cell Biol.* **155**, 937–948 (2001).
2. X Bi, RA Corpina, J Goldberg, Structure of the sec23/24–sar1 pre-budding complex of the copii vesicle coat. *Nature* **419**, 271–277 (2002).
3. J Jumper, et al., Highly accurate protein structure prediction with alphafold. *Nature* **596**, 583–589 (2021).
4. S Jo, T Kim, VG Iyer, W Im, Charmm-gui: a web-based graphical user interface for charmm. *J. Comput. Chem.* **29**, 1859–1865 (2008).
5. A Waterhouse, et al., Swiss-model: homology modelling of protein structures and complexes. *Nucleic Acids Res.* **46**, W296–W303 (2018).
6. W Humphrey, A Dalke, K Schulten, VMD – Visual Molecular Dynamics. *J Mol Graph* **14**, 33–38 (1996).
7. MJ Abraham, et al., Gromacs: High performance molecular simulations through multi-level parallelism from laptops to supercomputers. *SoftwareX* **1**, 19–25 (2015).
8. HJ Berendsen, D van der Spoel, R van Drunen, Gromacs: A message-passing parallel molecular dynamics implementation. *Comput. Phys. Commun.* **91**, 43–56 (1995).
9. J Huang, et al., Charmm36m: an improved force field for folded and intrinsically disordered proteins. *Nat. Methods* **14**, 71–73 (2017).
10. BR Brooks, et al., Charmm: the biomolecular simulation program. *J. Comput. Chem.* **30**, 1545–1614 (2009).
11. J Lee, et al., Charmm-gui input generator for namd, gromacs, amber, openmm, and charmm/openmm simulations using the charmm36 additive force field. *J. Chem. Theory Comput.* **12**, 405–413 (2016).
12. T Darden, D York, L Pedersen, Particle mesh ewald: An n log (n) method for ewald sums in large systems. *J. Chem. Phys.* **98**, 10089–10092 (1993).
13. WG Hoover, Canonical dynamics: Equilibrium phase-space distributions. *Phys. Rev. A* **31**, 1695 (1985).
14. S Nosé, A molecular dynamics method for simulations in the canonical ensemble. *Mol. Phys.* **52**, 255–268 (1984).
15. M Parrinello, A Rahman, Polymorphic transitions in single crystals: A new molecular dynamics method. *J. Appl. Phys.* **52**, 7182–7190 (1981).
16. B Hess, H Bekker, HJ Berendsen, JG Fraaije, Lincs: a linear constraint solver for molecular simulations. *J. Comput. Chem.* **18**, 1463–1472 (1997).
17. S Paul, SRK Ainavarapu, R Venkatramani, Variance of atomic coordinates as a dynamical metric to distinguish proteins and protein–protein interactions in molecular dynamics simulations. *J. Phys. Chem. B* **124**, 4247–4262 (2020).
18. YZ Ohkubo, TV Pogorelov, MJ Arcario, GA Christensen, E Tajkhorshid, Accelerating membrane insertion of peripheral proteins with a novel membrane mimetic model. *Biophys. J.* **102**, 2130–2139 (2012).
19. MG Hanna, et al., Sar1 gtpase activity is regulated by membrane curvature. *J Biol Chem* **291**, 1014–1027 (2016).
20. S Jo, JB Lim, JB Klauda, W Im, Charmm-gui membrane builder for mixed bilayers and its application to yeast membranes. *Biophys. J.* **97**, 50–58 (2009).
21. Y Qi, et al., Charmm-gui hmmm builder for membrane simulations with the highly mobile membrane-mimetic model. *Biophys. J.* **109**, 2012–2022 (2015).
22. HJ Berendsen, Jv Postma, WF Van Gunsteren, A DiNola, JR Haak, Molecular dynamics with coupling to an external bath. *J. Chem. Phys.* **81**, 3684–3690 (1984).
23. Z Wu, K Schulten, Synaptotagmin's role in neurotransmitter release likely involves ca2+-induced conformational transition. *Biophys. J.* **107**, 1156–1166 (2014).
